# Organization of the catecholaminergic system in the short-lived fish *Nothobranchius furzeri*

**DOI:** 10.1101/2021.06.30.450602

**Authors:** Janina Borgonovo, Patricio Ahumada-Galleguillos, Alejandro Oñate-Ponce, Camilo Allende-Castro, Pablo Henny, Miguel L. Concha

## Abstract

The catecholaminergic system has received much attention based on its regulatory role in a wide range of brain functions and its relevance in aging and neurodegenerative diseases. In the present study, we analyzed the neuroanatomical distribution of catecholaminergic neurons based on tyrosine hydroxylase (TH) immunoreactivity in the brain of adult *Nothobranchius furzeri*. In the telencephalon, numerous TH+ neurons were observed in the olfactory bulbs and the ventral telencephalic area, arranged as strips extending through the rostrocaudal axis. We found the largest TH+ groups in the diencephalon at the preoptic region level, the ventral thalamus, the pretectal region, the posterior tuberculum, and the caudal hypothalamus. In the dorsal mesencephalic tegmentum, we identified a particular catecholaminergic group. The rostral rhombencephalon housed TH+ cells in the locus coeruleus and the medulla oblongata, distributing in a region dorsal to the inferior reticular formation, the vagal lobe, and the area postrema. Finally, scattered TH+ neurons were present in the ventral spinal cord and the retina. From a comparative perspective, the overall organization of catecholaminergic neurons is consistent with the general pattern reported for other teleosts. However, *Nothobranchius furzeri* shows some particular features, including the presence of catecholaminergic cells in the midbrain. This work provides a detailed neuroanatomical map of the catecholaminergic system of *Nothobranchius furzeri*, a powerful aging model, also contributing to the phylogenetic understanding of one of the most ancient neurochemical systems.

## 1 Introduction

Historically, the catecholaminergic (CAergic) system has aroused particular interest due to its role in a variety of neural functions, including motor control, memory, learning, reward, motivation, sleep, temperature regulation, and reproduction (Crocker, 1997; Hurley et al., 2004; Salamone and Correa, 2012). The CAergic system has also attracted attention from the clinical perspective since imbalances in catecholamines or components of the CAergic system are present in neuropsychiatric disorders such as schizophrenia, depression, and anxiety (Davis et al., 1994; Joyce, 1993), and in neurodegenerative diseases with high prevalence as is the case of Parkinson’s disease (PD) (Dawson and Dawson, 2003). In PD, the dopaminergic (DAergic) neurons of the substantia nigra (SN) are severely affected, and other CAergic cells also suffer changes during the disease’s development. For instance, TH+ neurons increase in the olfactory bulb of PD patients (Huisman et al., 2004, 2008) and the striatum of animal models of PD (El Massri et al., 2017; Huot et al., 2007; Palfi et al., 2002; Tande et al., 2006; Tashiro et al., 1990; Tashiro et al., 1989). Moreover, the CAergic system deteriorates in the course of physiological aging, showing an age-dependent loss of neuronal components in the locus coeruleus (Manaye et al., 1995), striatum (Huot et al., 2007) and retina (Roufail and Rees, 1997). Together, these findings reveal that the CAergic system’s deterioration is a cross-cutting issue of normal and pathological aging.

The organization of the CAergic system has been reported in a wide range of vertebrates, from cyclostomes to mammals, since the first demonstration of catecholamines in the central nervous system (CNS) (Bertler and Rosengren, 1959; Carlsson, 1959; Montagu, 1956; von, 1961). These studies have revealed an overtly conserved distribution of CAergic neuronal groups throughout evolution (Smeets and Gonzalez, 2000). Such conservation has especially been relevant for biomedical research since it allowed drawing parallels between humans and species of biomedical relevance, such as mouse and some teleost fish species, in the search for models of neurodegenerative disorders. Among the latter, zebrafish and medaka have gained space as key non-mammalian vertebrate models for studying neurological diseases and PD (Matsui and Takahashi, 2018; Matsui et al., 2014; Wang et al., 2017). These teleost species show a complex CAergic system, containing most of the CAergic groups found in mammals based on anatomical, molecular and connectivity criteria (Filippi et al., 2012; Rink and Wullimann, 2001; Tay et al., 2011). Furthermore, the TH+ cells found in the posterior tuberculum of these species have been considered analogous to the mammalian SN and, as seen in mammals, these neurons are especially vulnerable to toxic insults and the expression of mutant proteins linked to PD (Sallinen et al., 2009; Sheng et al., 2010; Soman et al., 2019).

In the last decade, the turquoise killifish (*Nothobranchius furzeri*) has reached popularity as a model organism for aging research (Genade et al., 2005; Terzibasi et al., 2009). *Nothobranchius furzeri* belongs to the order Cyprinodontiformes and inhabits temporary puddles in the African savannah, showing a life cycle adapted to the temporary habitat in which fish live. In a short period of the year that coincides with the rainy season, fish hatch from the eggs, reach sexual maturity and reproduce. When the puddles dry, the adults die. Still, the eggs remain buried in the mud resisting desiccation until the environmental conditions are adequate to restart the life cycle (Blazek et al., 2013; Reichard et al., 2009). Since introducing the first individuals of the Gona Re Zhou (GRZ) strain into the laboratory, the use of *Nothobranchius furzeri* has persistently grown due to their short lifespan and accelerated aging, in which fish express biological markers of aging common to other vertebrates, including humans (Baumgart et al., 2015; Di Cicco et al., 2011; Genade et al., 2005; Terzibasi et al., 2009; Valenzano et al., 2006). Recent work also showed that *Nothobranchius furzeri* develops a PD-like pathology in an age-dependent manner with physiological loss of the TH+ cells in the posterior tuberculum, deposition of α-synuclein aggregates, and motor disability (Matsui et al., 2019). Together, the short lifespan, accelerated aging and age-related loss of CAergic neurons make *Nothobranchius furzeri* a versatile and tractable model for longitudinal studies with a biomedical perspective, especially for pathologies associated with age like PD. However, although information of the CAergic system is available for zebrafish and many other teleosts (Bhat and Ganesh, 2017; Brinon et al., 1998; Goebrecht et al., 2014; Hornby and Piekut, 1990; Manso et al., 1993; Meek and Joosten, 1993; Parent and Northcutt, 1982; Roberts et al., 1989; Rodriguez-Gomez et al., 2000), a systematic description of the different neuronal groups that compose this system has not been performed in *Nothobranchius furzeri*. This missing information is relevant since the CAergic system is heterogeneous in its composition, and knowledge of the diverse neuronal types is essential, considering that they are differentially affected during physiological aging and under pathological conditions (Huot et al., 2007; Manaye et al., 1995; Roufail and Rees, 1997).

The present study aims to provide a comprehensive analysis of the CAergic system’s anatomy in *Nothobranchius furzeri*. For this, we used tyrosine hydroxylase (TH) immunofluorescence to locate the CAergic cells and fibers in the turquoise killifish brain. We focused on describing the spatial organization of the different neuronal groups and the morphological features of TH+ cells. Our results show that the overall organization of the CAergic system of *Nothobranchius furzeri* is comparable to other teleosts. However, killifish shows a distinct TH+ group in the dorsal midbrain that resembles CAergic neuronal groups observed in some species of holosteans and cladistians. This work provides a detailed neuroanatomical framework for aging studies related to the CAergic system in *Nothobranchius furzeri*, and contributes to our understanding of the evolution of the CAergic system in vertebrates, especially of the mesencephalic groups.

## 2 Materials and Methods

### 2.1 Fish maintenance and husbandry

Adult wild type *Nothobranchius furzeri* of the Gona Re Zhou (GRZ) strain were raised and maintained in the fish facility of the Laboratory of Experimental Ontogeny, from the Faculty of Medicine, University of Chile. Fish were kept under 12/12 h light/dark cycle regime at 26°C, in a recirculating system. The conductivity of water was set to 400-500 μS and the pH to 7–7.5. Adult fish were kept at density of one male or 3 females per tank. The food was supplied 3 times a day with newly hatched brine shrimps and once with freshwater live food (*Lumbricus variegatus*). Weekly, 50% of total water volume was replaced with fresh water to remove the waste produced by fish. All animal procedures were approved by the Bioethics Committee of the Faculty of Medicine, University of Chile (CICUA certificate number: 20385-MED-UCH).

### 2.2 Tissue sampling

Experiments were performed on *Nothobranchius furzeri* brains of both sexes from 2 to 4 months old. The total of animals used for the experiments were 14. Individuals were euthanized with a 0.1% solution of ethyl 3-aminobenzoate methane sulfonate (MS-222; Sigma, St. Louis, MO, USA; A-5040). The brains were removed and fixed by immersion in 4% w/v paraformaldehyde in 0.1 M phosphate buffer (PB, pH 7,2 at 4 °C) for 24 h and subsequently were embedded in 2% low melting-agarose. The tissue was cut in coronal, sagittal or horizontal orientation at 150-200 µm of thick using the vibroslice (Campden MA752). The brain sections were collected in PBS 1X.

### 2.3 Immunofluorescence

The immunofluorescence was performed according to Matsui and Sugie (2017). In brief, the floating slices were incubated in 10 mM sodium citrate buffer, pH 8.5, at 80°C for 120 min, washed three times in wash buffer (PBS, Triton X-100 1%) and blocked for 30 min in blocking buffer (PBS, Triton X-100 1%, 2% BSA). Then, the slices were incubated with anti-tyrosine hydroxylase (Millipore Cat# AB152), anti-HuC/HuD (Thermo Fisher Scientific Cat# A-21271) or anti-acetylated α-tubulin (Sigma-Aldrich Cat# T6793) primary antibody overnight at 4°C, all at dilution of 1/200. After washing, the sections were incubated with the respective secondary antibody and counterstained with hoechst 33258 (1:1000, Thermo Fisher Scientific Cat# H3569) overnight at 4°C. The secondary antibodies used were anti-rabbit Alexa Fluor 488 (1:200, Molecular Probes Cat#A11070) and anti-mouse Alexa Fluor 568 (1:200, Invitrogen Cat#A11004). Lastly, the tissue was washed three times in washing buffer and the sections mounted with Vectashield (Vector Cat#VC-H-1400-L010) on glass slides for imaging.

The specificity of the TH antibody used in the present study was previously verified by Matsui et al. (2019). In this study, different controls were performed in brain tissue of adult *Nothobranchius furzeri*, including western blot analysis and comparison of TH immunostaining with the expression pattern of the dopamine transporter (*dat*) and the noradrenaline transporter (*net*) obtained after *in situ* hybridization (Matsui et al., 2019).

### 2.4 Imaging

The immunofluorescent images were acquired using a Leica TCS LSI macro zoom confocal microscope with a 5x objective. In addition, to obtain a cellular resolution, the slices were imaged with a Volocity ViewVox spinning disc (Perkin Elmer) coupled to a Zeiss Axiovert 200 inverted microscope using a Plan-Apochromat 40x/1.2W or 10x/0.3W objective with a laser 488/520 (excitation/emission wavelengths). To determine the cell size, we measured the largest diameter of the soma in the selected neurons. The processing and analysis of digital images including measurement of soma diameter were performed using ImageJ (http://rsb-web.nih.gov/ij/) and Adobe Photoshop CS3 (Adobe).

## 3 Results

The main subdivisions of the brain, the localization of brain nuclei, and the neuroanatomical terminology used in this paper is based on D’Angelo (2013), the only available brain atlas of adult *Nothobranchius furzeri*. For the caudal sections of the rhombencephalon we included information from the brain atlas of zebrafish developed by Wullimann et al. (1996). We followed the classical forebrain subdivision into telencephalon and diencephalon consistent with D’Angelo (2013), where the diencephalon includes the pretectal area, thalamus, hypothalamus, and preoptic areas. However, it is important to bear in mind that this classical paradigm differs from the prosomeric model proposed by Puelles and Rubenstein (2003), where the forebrain is divided into three caudorostral segments (p1, p2 and p3), and a secondary prosencephalon. Within this frame, the pretectum, thalamus (dorsal thalamus), and prethalamus (ventral thalamus) are the alar part of the p1, p2, and p3 prosomeres respectively, while the hypothalamic and preoptic areas are part of the secondary prosencephalon together with the telencephalon (Puelles and Rubenstein 2003).

The commercially available TH antibodies only detect the product of *th1*, one of the two tyrosine hydroxylase genes (*th1* and *th2*) present in non-mammalian vertebrates. However, most CAergic cells in non-mammalian vertebrate species express *th1*. In contrast, *th2* is found only in the cerebrospinal fluid-contacting cells (CSF-c) of the preoptic area and hypothalamic regions (Yamamoto et al., 2010). Thus, in this study, we present the neuronal groups that probably express the *th1* transcript. Henceforth, we name these TH-immunoreactive cells as TH+.

### 3.1 Telencephalon

#### Olfactory bulbs

The olfactory bulbs (OB) of *Nothobranchius furzeri* are located in the ventral part of the rostralmost telencephalic hemispheres. These regions showed a large population of TH+ cells distributed predominantly at the outer zone, along the dorsal region of the external cell layer (ECL) and less abundantly in the internal cell layer (ICL). Conspicuous TH+ fibers were present in the glomerular layer (GL) (**Figures 1A; 2A-B; S1A; S3C**). TH+ neurons were ovoid in shape varying in size from 10 to 12 µm. A single process emerged from the cell bodies and TH+ fibers were generally oriented towards the internal area (**Figure 7A**).

**Figure 1.**
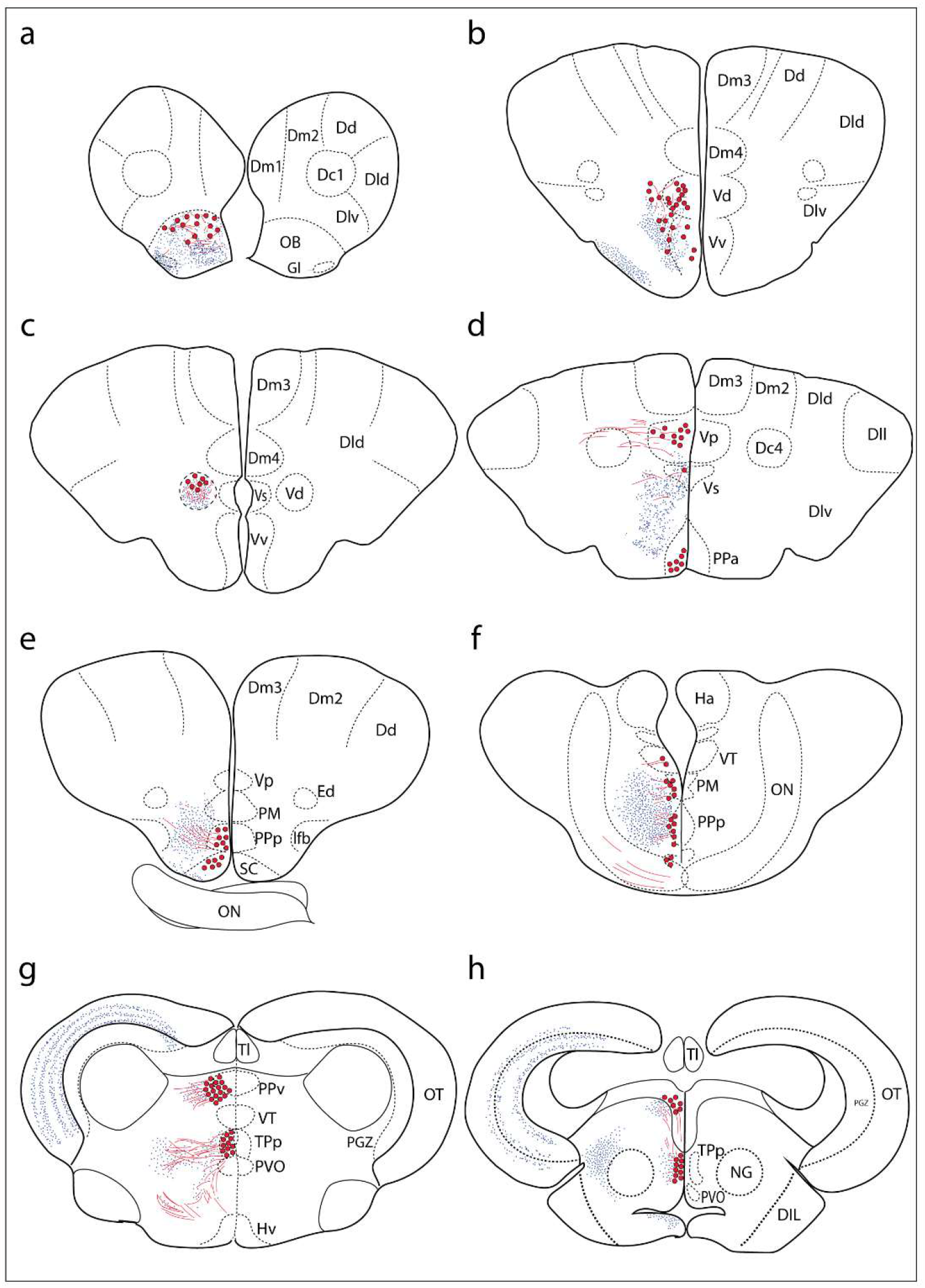
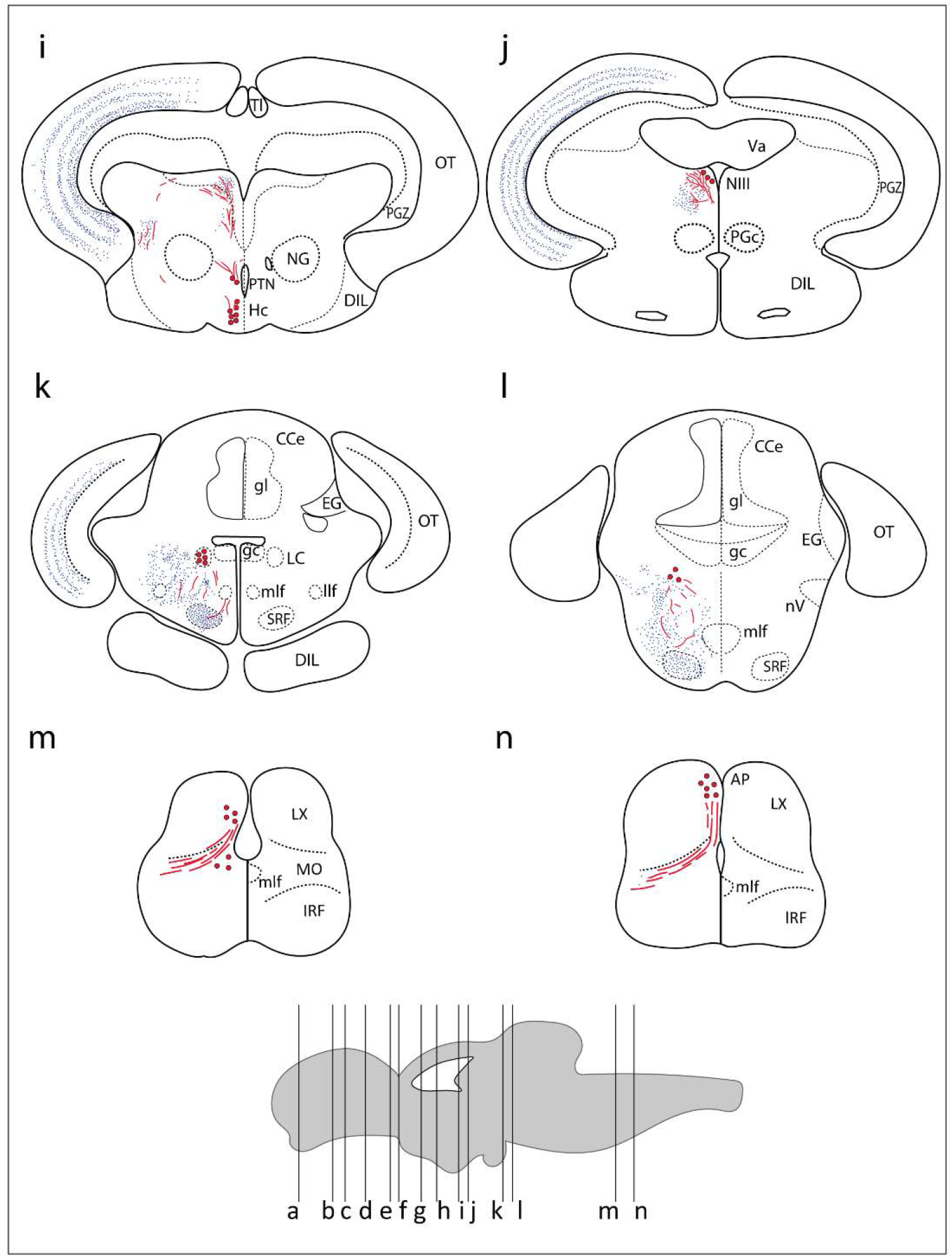
Summary of TH immunoreactivity in the brain of *Nothobranchius furzeri*. **(A-N)** Diagrams of transverse sections showing TH+ cells bodies (large red dots), fibers (wavy red lines) and nerve terminals (small blue dots). The rostrocaudal level of each section is indicated in the lower brain scheme. See list of abbreviations, which applies to all figures.

**Figure 2.**
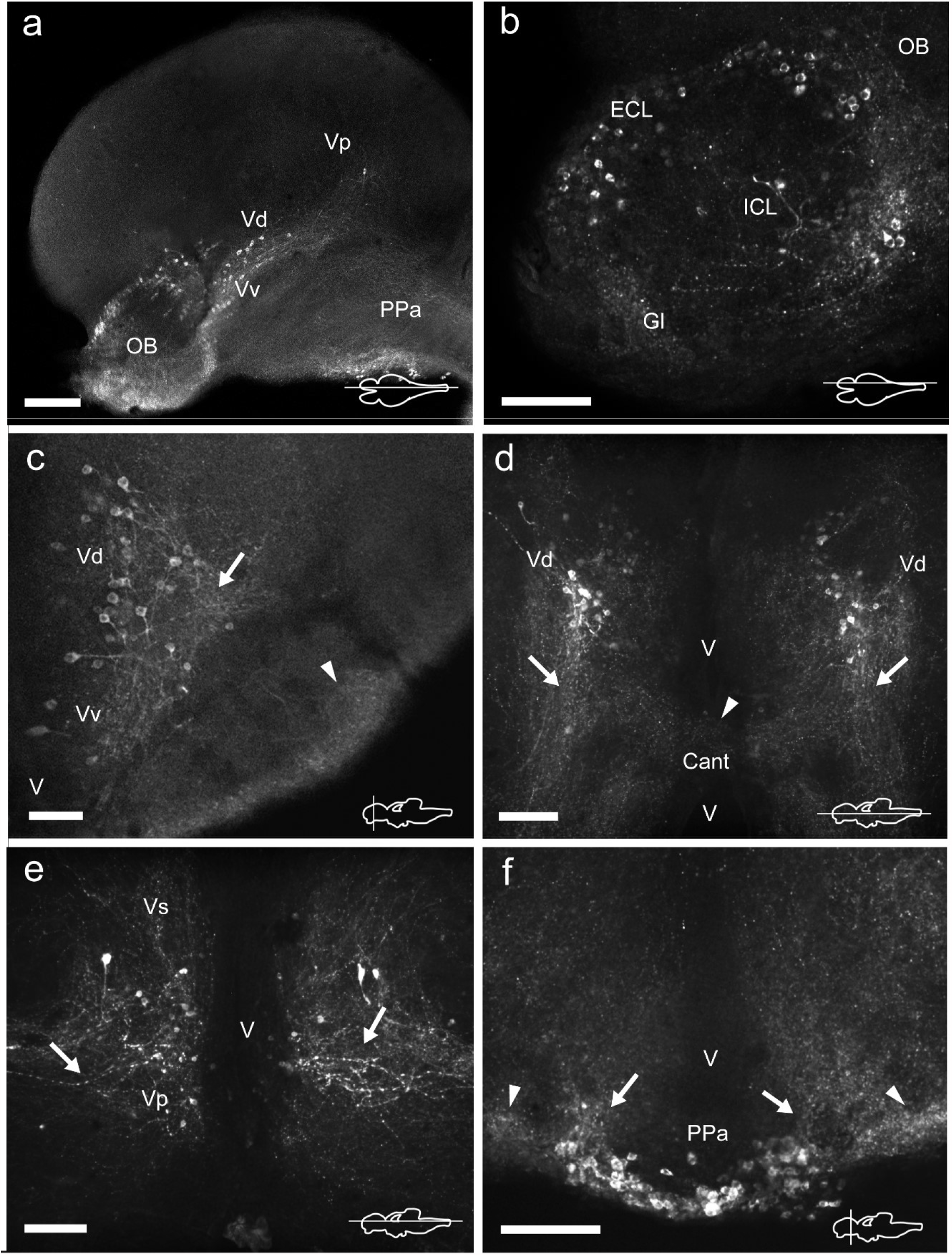
TH immunoreactive neurons and fibers in the telencephalon and anterior preoptic area of *Nothobranchius furzeri*. **(A)** Para-sagittal section showing a panoramic view of TH+ neuronal groups in the olfactory bulb, subpallium and in the anterior preoptic nucleus. **(B)** Para-sagittal section showing the TH+ groups in the olfactory bulb. **(C-E)** Ventral telencephalic areas showing TH+ groups in the ventral, dorsal, supracomissural and posterior zones. **(C)** Transverse section showing local TH+ processes of ventral/dorsal zones (arrow) and intense immunoreactive innervation at the ventrolateral edge of the telencephalon (arrowheads). **(D)** Horizontal section showing the caudalward projections of the dorsal zone (arrows) and the TH+ fibers at the level of the anterior commissure (arrowhead). **(E)** Horizontal section showing the lateral processes of the posterior zone (arrows). **(F)** Transverse section showing the TH+ neurons at the anterior preoptic nucleus. Arrows and arrowheads indicate the TH+ dorsocaudal projections of the anterior preoptic neurons and the lateral intense neuropil seen at each side of this cell group, respectively. For abbreviations see list. The orientation/level of sections is shown at the bottom right corner of each panel. Dorsal is to the top in transverse sections. Rostral is to the top and left, in horizontal and para-sagittal sections, respectively. Scale bars, 250µm (a) and 100 µm (b-f).

#### Telencephalic hemispheres

The telencephalic hemispheres of *Nothobranchius furzeri*, as in other vertebrates, are subdivided into pallial (dorsal) and subpallial (ventral) regions. While the pallium was devoid of TH+ cells, the subpallium harbored populations of CAergic neurons in its ventral (Vv), dorsal (Vd), supracomissural (Vs) and posterior (Vp) nuclei (**Figure 1B-D**). These TH+ neurons were disposed as a continuous strip, extending rostrocaudally from the caudal end of the OB to the Vp (**Figures 2A; S2A**). The neurons of the rostral regions projected locally and also sent fibers in a rostroventral direction (**Figures 2C; S1B-C**). The most caudal TH+ cells of the Vd also projected caudalward and some fibers crossed the anterior commissure (**Figures 2D; S3B**). In turn, TH+ neurons in the Vs and Vp presented long laterocaudal processes (**Figures 2E; S1D, S3A**). TH+ cells of all these areas shared a similar morphology being round in shape with medium immunoreactivity level and showing an average size of 11-15 µm (**Figure 7B**).

### 3.2 Diencephalon

The diencephalon of *Nothobranchius furzeri* showed the largest number of TH+ groups, which were located in the following regions: preoptic area (PO), thalamus, pretectum, posterior tuberculum (TP) and hypothalamus.

#### Preoptic area

A group of TH+ cells was present in the anterior parvocellular preoptic nucleus (PPa), caudal to the anterior commissure (Cant) and at the floor of the telencephalon (**Figures 1D; 2A,F; S1D; S2A-C**). These neurons elaborated lateral projections that ended in an intensely labelled neuropil at each side of the PPa (**Figures 2F; S1D**). In addition, PPa neurons emitted long processes dorsocaudally, which fasciculated to form the preopticohypothalamic tract (poht) (**Figures 2F; S2A-C**). The PPa was constituted by small round cells with an average size of 8-10 µm and intense TH immunoreactivity (**Figure 7C**). In a position caudal to the PPa, the suprachiasmatic nucleus (SC) contained a few scattered TH+ cells (**Figures 1E; S1E; S2B-C**). The SC neurons showed a similar morphology to those of the PPa and their projections joined the poht (**Figure S2B-C**). Further, two groups of TH+ cells were observed along the posterior parvocellular preoptic nucleus (PPp) and magnocellular preoptic nucleus (PM) (**Figures 1E-F; 3A-B,E-F; S1E-F; S2B-C**). The neurons of both nuclei were located against the ventricular surface and their round somas exhibited a diameter of 11-14µm (**Figure 7E**). PM neurons showed intense TH immunoreactivity and displayed a radial arborization, which extended laterally to the adjacent PO regions. In contrast, PPp neurons were lightly immunostained and their processes were not discernible (**Figure 7E**).

#### Epithalamus, thalamus and pretectal region

The epithalamus lacked CAergic cells, however the habenula (Ha) showed discrete TH+ innervation (**Figure S3A**). In the thalamus, at the ventral subdivision (VT) (also called prethalamus), a few TH+ cells were found away from the ventricular zone (**Figures 3A-B,D; S1E; S2B-C**). Their lateral processes extended toward the lateral edge of the diencephalon. These neurons were weakly stained and measured less than 11 µm (**Figure 7F**).

**Figure 3.**
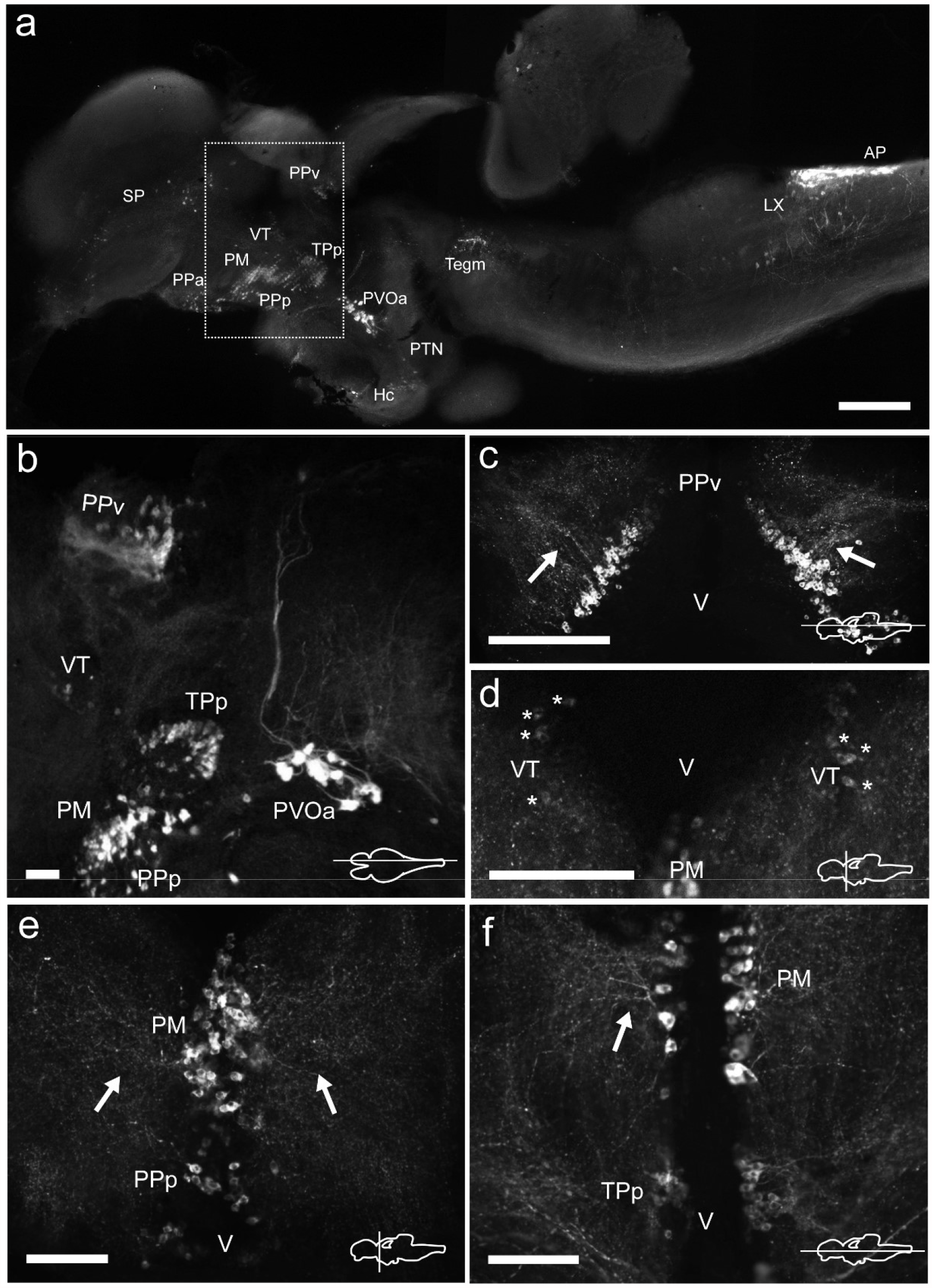
TH immunoreactive neurons and fibers in the diencephalon of *Nothobranchius furzeri* at the preoptic, thalamic and pretectal levels. **(A)** Para-sagittal section of the brain showing a panoramic distribution of TH+ groups. The dotted box delimits the neuronal groups analyzed in this figure. **(B)** Para-sagittal section showing the TH+ groups at the ventral periventricular pretectal nucleus, ventral thalamic nucleus, parvocellular portion of preoptic nucleus, magnocellular preoptic nucleus and paraventricular organ-accompanying cells. **(C)** Horizontal section showing the densely packed TH+ neurons of the ventral periventricular pretectal nucleus sending projections towards to the optic tectum (arrowheads). **(D)** Transverse section showing TH+ neurons at the ventral thalamus with characteristic slightly stained somas (arrows). **(E)** TH+ cells of the parvocellular portion of preoptic nucleus and the magnocellular preoptic nucleus with their lateral projections as seen in a transverse section (arrows). **(F)** Horizontal section showing TH+ neurons and their projections (arrowheads) at the magnocellular nucleus and its relative position with respect to the periventricular nucleus of the posterior tuberculum. For abbreviations see list. The orientation/level of sections is shown at the bottom right corner of each panel. Dorsal is to the top in transverse sections. Rostral is to the top and left in horizontal and para-sagittal sections, respectively. Scale bars, 500 µm (a) and 100 µm (b-f).

A densely-packed population of TH+ neurons was observed at the pretectal nucleus, in the ventral periventricular subdivision (PPv). Their TH+ fibers followed a dorso-lateral trajectory to reach the rostral region of the optic tectum (OT) (**Figures 1G; 3A-C; S1G; S2A-C; S3A**). The pretectal cells exhibited intense immunoreactivity and round cell bodies with an average some size of 8-12 µm (**Figure 7D**).

#### Posterior tuberculum and hypothalamus

Three groups of TH+ cells were present in the posterior tuberculum. The rostralmost group was localized in the periventricular nucleus of the posterior tuberculum (TPp), positioned ventral to the VT (**Figures 1G; S2B-C; S3C**). A tuft of processes emerged laterally from the nucleus and ended in the vicinity (**Figures 4A-B,D; S1G**). TPp neurons presented moderate TH immunostaining, were densely packed and showed round soma with a diameter of 10-13 µm (**Figure 7G**). The second group appeared in a more ventral and caudal position to the TPp. This group was not strictly associated with a specific nucleus, but rather it extended in the boundaries of the TPp and the paraventricular organ (PVO), and we decided to name it as paraventricular organ-accompanying cells of the posterior tuberculum (PVOa), as others authors (**Figures 1H; S1H; S2A-C; S3C**) (Ma, 2003). The PVOa consisted of a few large, multipolar and intensely labeled neurons (**Figure 4A,C,D**). We could distinguish two neuronal subpopulations within the PVOa. The first adopted a more rostral position and the neurons showed a soma diameter of 20-25 µm (**Figure 7O**). These cells sent axonal projections in rostroventral and lateral directions towards the telencephalon and hypothalamus (**Figures 4A; S2A-C; S3C**). The second PVOa group appeared more caudally, their cells bodies were slightly smaller with a diameter of 14-18 µm and emitted thick processes with dorsal orientation (**Figure 7P**). These axons travelled large distances to reach the floor of the tectal ventricle (TeV) where they turned 90° caudalward to descend towards the medulla (**Figures 4A; 5A-B; S2A-C; S3C**). The third TH+ group of the posterior tuberculum was observed in the posterior tuberal nucleus (PTN) (**Figures 1I; 4A,E; S1I; S2B-C**). It was composed of a few teardrop shape neurons scattered on each side of the ventricle with ascending processes that extended towards the PVOa (**Figure 7H**).

**Figure 4.**
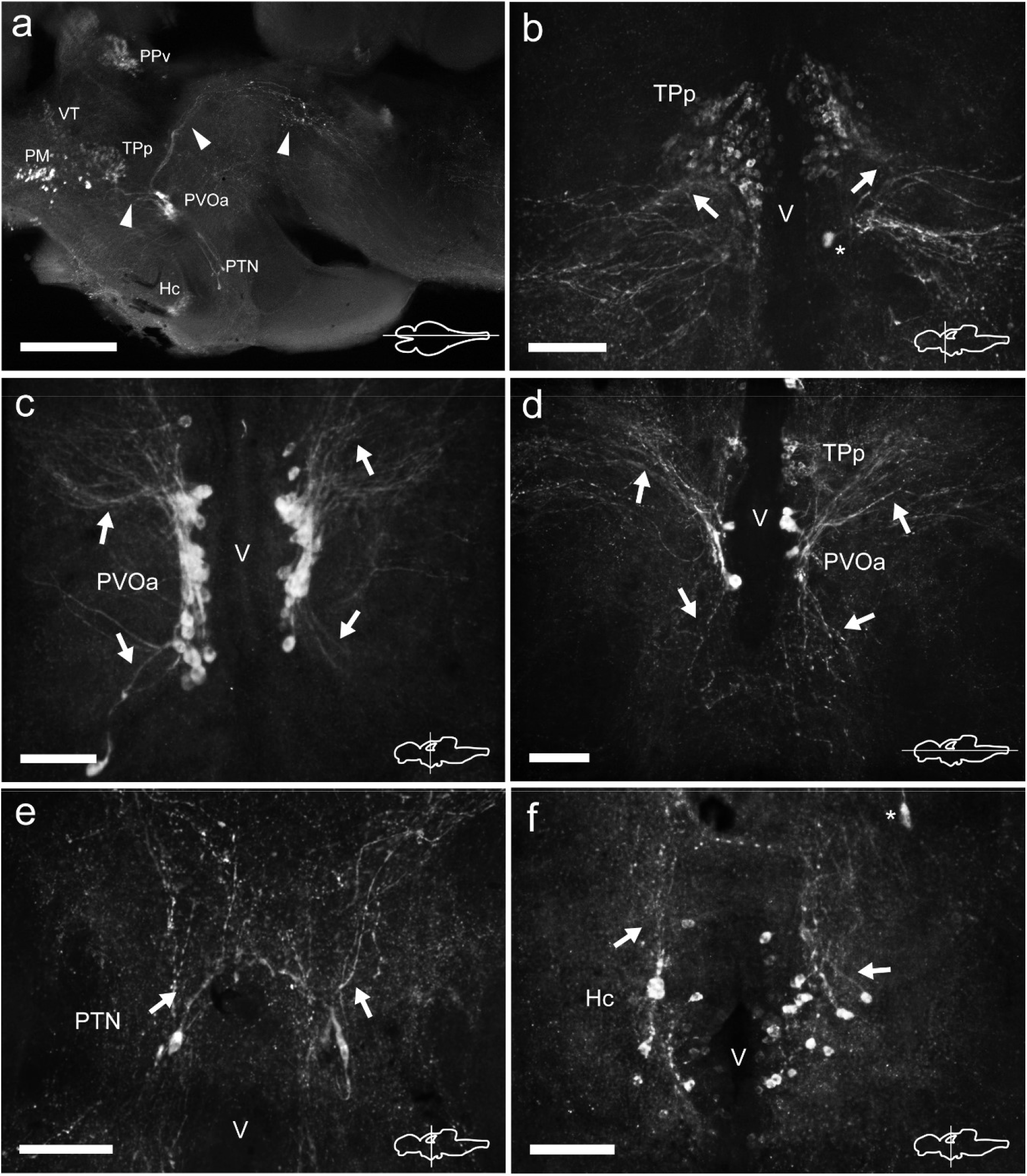
TH immunoreactive neurons and fibers in the diencephalon of *Nothobranchius furzeri* at the level of the posterior tuberculum and hypothalamus. **(A)** Para-sagittal section showing a panoramic view of the distribution of all TH+ groups in the diencephalon. Arrowheads indicate TH+ fibers travelling from the paraventricular organ-accompanying cells towards the telencephalon and hypothalamus (bottom left), and across the mesencephalon to turn posteriorly towards the medulla (top right). **(B)** Transverse section showing the TH+ group in the periventricular nucleus of the posterior tuberculum. Arrowheads show the lateral tuft of projections emerging from the nucleus. The arrow indicates a soma of the paraventricular organ-accompanying cells. **(C)** Transverse section showing the large pear shape neurons of the TH+ paraventricular organ-accompanying cells group. Arrowheads indicate the ascending and descending processes. **(D)** Horizontal section showing the relative position of the periventricular nucleus of the posterior tuberculum and the paraventricular organ-accompanying cells. Arrowheads point to ascending/descending processes of the paraventricular organ-accompanying cells. **(E)** Transverse section showing TH+ neurons at the posterior tuberal nucleus. The few cells of this group are positioned at each side of the ventricle and send projections in the dorsal direction (arrowheads). **(F)** Transverse section showing the TH+ neurons at the caudal hypothalamus. The arrowhead indicates the orientation of processes. The arrow indicates a cell of the posterior tuberal nucleus. For abbreviations see list. The orientation/level of sections is shown at the bottom right corner of each panel. Dorsal is to the top in transverse sections. Rostral is to the top and left, in horizontal and para-sagittal sections, respectively. Scale bars, 500 µm (a) and 100 µm (b-f).

At the hypothalamus, TH+ neurons were observed in the caudal subdivision (Hc), dorsal to the posterior recess (rec) (**Figures 1I; 4A,F; S1I,J; S2A-C**). The cell bodies were small and round with a size of 9-12 µm and showed dorsally and ventrolaterally oriented processes that projected locally in the adjacent brain regions (**Figure 7I**). The hypothalamus also showed a dense network of TH+ fibers on the ventral surface, originated from the preoptic area and the ventral diencephalon (**Figure S2A-C**).

### 3.3 Mesencephalon

The dorsal mesencephalon of *Nothobranchius furzeri* is composed of two large optic tectum (OT) and the torus longitudinalis (Tl). Both structures lacked CAergic cells, but the OT showed a dense innervation, with TH+ fibers splaying out along the tectal layers (**Figures 1G-K; S1G-L; S3A-C**).

In the midbrain tegmentum, the torus semicircularis (TS) also presented TH+ neuropil, especially in the periventricular and lateral zones (**Figures 1J; S1K; S3A-B**). In addition, intense and varicose CAergic fibers were observed in the tegmentum at the level of the oculomotor nucleus (NIII). Many of these fibers originated from the TP (**Figures 1J; 5A-B; S1K; S2A-C**). Interestingly, a small group of TH+ positive neurons were located close to the ventricle and dorsal to the oculomotor nucleus in the mesencephalic tegmentum (tegm), where a TH+ plexus was placed (**Figures 1J; 5A; S2C**). These cells showed weak TH immunofluorescence and presented a pear shape morphology (**Figure 7J**).

### 3.4 Rhombencephalon

#### Cerebellum and rostral rhombencephalon

The cerebellum of *Nothobranchius furzeri* showed low density of TH+ fibers in the corpus of the cerebellum (CCe) and in the granular eminence (EG) (**Figure S1L-M**).

In the rostral rhombencephalon, a small group of TH+ cells were found dorsal to the superior reticular formation (SRF) (**Figures 1K-L; 5B-C; S1L-M; S2B; S3A**). This group corresponded to the locus coeruleus (LC) and was composed of large pear shape neurons with an average size of 25-30 µm, showing long branched processes directed ventrally (**Figure 7K**).

#### Medulla oblongata

At the caudal rhomencephalic level, a longitudinal column of CAergic neurons was observed in rostrocaudal direction from the caudal end of LVII to the medullospinal junction (**Figure S2A,C**). This column consisted of two neuronal populations distinguishable by their morphology. The rostral group (MO) was positioned in a position dorsomedial to the intermediate (IMRF) and inferior (IRF) reticular formation (**Figures 1M; 5D-E**). These cells showed a multipolar shape and presented lateroventral processes (**Figure 7L**). The second group was positioned more caudally, in the vagal lobe (LX) and consisted of teardrop shape neurons with an intense TH immunoreactivity and lateroventral processes that reached the lateral edge of the medulla (**Figures 1M; 5B,D,F-G; 7M; S1O-Q; S2A-C**). An additional TH+ group was present in the medulla oblongata, at the level of the area postrema (AP) (**Figures 1N; 5D,G; S1P-Q; S2C; S3A-B**), where a densely packed group of the round cells formed a wedge in the midline (**Figure 7N**).

Many regions of the medulla oblongata showed CAergic innervation. The medial octavolateralis nucleus (MON) had an intense TH+ neuropil and the SRF/IMRF/IRF also presented a moderate innervation, which was more prominent in the ventral surface (**Figures S1M-P; S2A-C**). Besides, a longitudinal tract (lt), probably originated in the diencephalic area, spanned all the rostrocaudal extension of the rhombencephalon to reach the spinal cord (**Figures S1O-Q; S2A-B**).

### 3.5 Spinal Cord

The spinal cord (sc) showed a reduced number of small lightly immunostained CAergic neurons in the ventral aspect (**Figure 5H**), away from the central canal. In addition, many TH+ fibers coursed in rostrocaudal direction, predominantly at the medial and ventral positions of the sc (**Figure 5H**).

**Figure 5.**
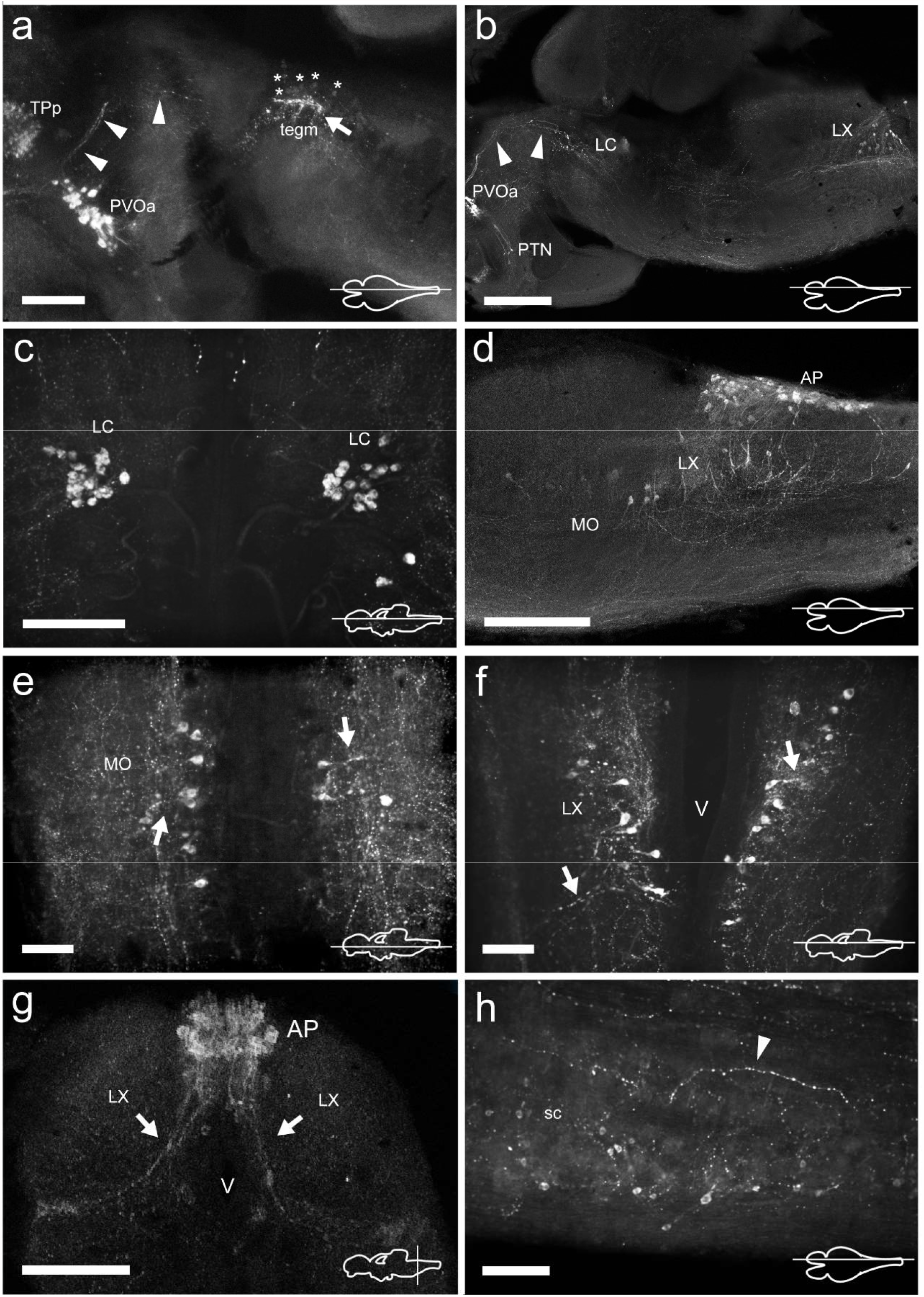
TH immunoreactive neurons and fibers in the mesencephalon, rhombencephalon and spinal cord of *Nothobranchius furzeri*. **(A)** Para-sagittal section showing the group of mesencephalic TH+ neurons at the level of the oculomotor nucleus (asterisks indicate their lightly stained somas) and the intense TH+ plexus (arrow). TH+ fibers travelling from the paraventricular organ-accompanying cells across the mesencephalon to turn posteriorly towards the medulla are indicated with arrowheads. **(B)** Para-sagittal section along the extension of the rhombencephalon showing the most rostral TH+ group at the locus coeruleus and the caudal group in the vagal lobe. Arrowheads point to the same TH+ projections as in (a). **(C)** Horizontal section showing the TH+ group of the locus coeruleus. **(D)** Para-sagittal section showing the distribution of caudal rhombencephalic TH+ groups (medulla oblongata, vagal lobe and area postrema). **(E-F)** Horizontal sections showing the TH+ groups at the medulla oblongata and vagal lobe. Arrows indicate the lateral projections. **(G)** Transverse section showing the densely packed group of TH+ neurons at the area postrema. Arrows indicate the ventral processes. **(H)** Para-sagittal section showing scattered small TH+ neurons in the spinal cord. The arrowhead points to a TH+ fiber in the ventral region of the spinal cord. For abbreviations see list. The orientation/level of sections is shown at the bottom right corner of each panel. Dorsal is to the top in transverse sections. Rostral is to the top and left, in horizontal and para-sagittal sections, respectively. Scale bars, 500 µm (a,d), 250µm (b,g) and 100 µm (c,e,f,h).

### 3.6 Retina

At the retina, a sparse population of TH+ neurons was observed in the inner nuclear layer (INL) and characteristically sent single process to the inner plexiform layer (IPL) (**Figure 6**). Based on their general morphology, localization and orientation we suggest these neurons correspond to amacrine cells.

**Figure 6.**
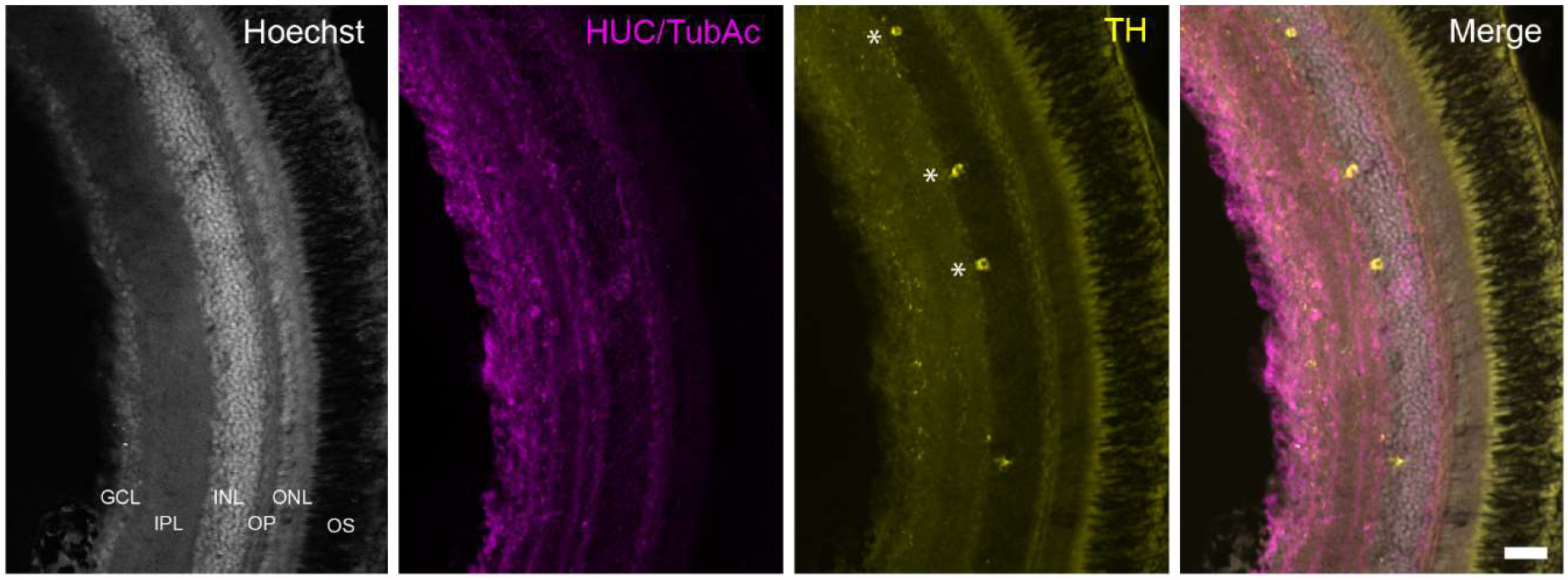
TH+ cells in the retina of *Nothobranchius furzeri*. **(A)** Hoechst stain showing the basic organization of the retina into the ganglion cell layer, inner plexiform layer, inner nuclear layer, outer plexiform layer, outer nuclear layer and outer segment. **(B)** HUC/TubAc immunohistochemistry (magenta) showing the spatial distribution of somas and axons within the retina layers. **(C)** TH immunohistochemistry (yellow). Asterisks indicate TH+ amacrine cells. **(D)** Merge of the staining in a-c showing the localization of TH+ cells in the inner nuclear layer. For abbreviations see list. Scale bar, 100 µm.

**Figure 7.**
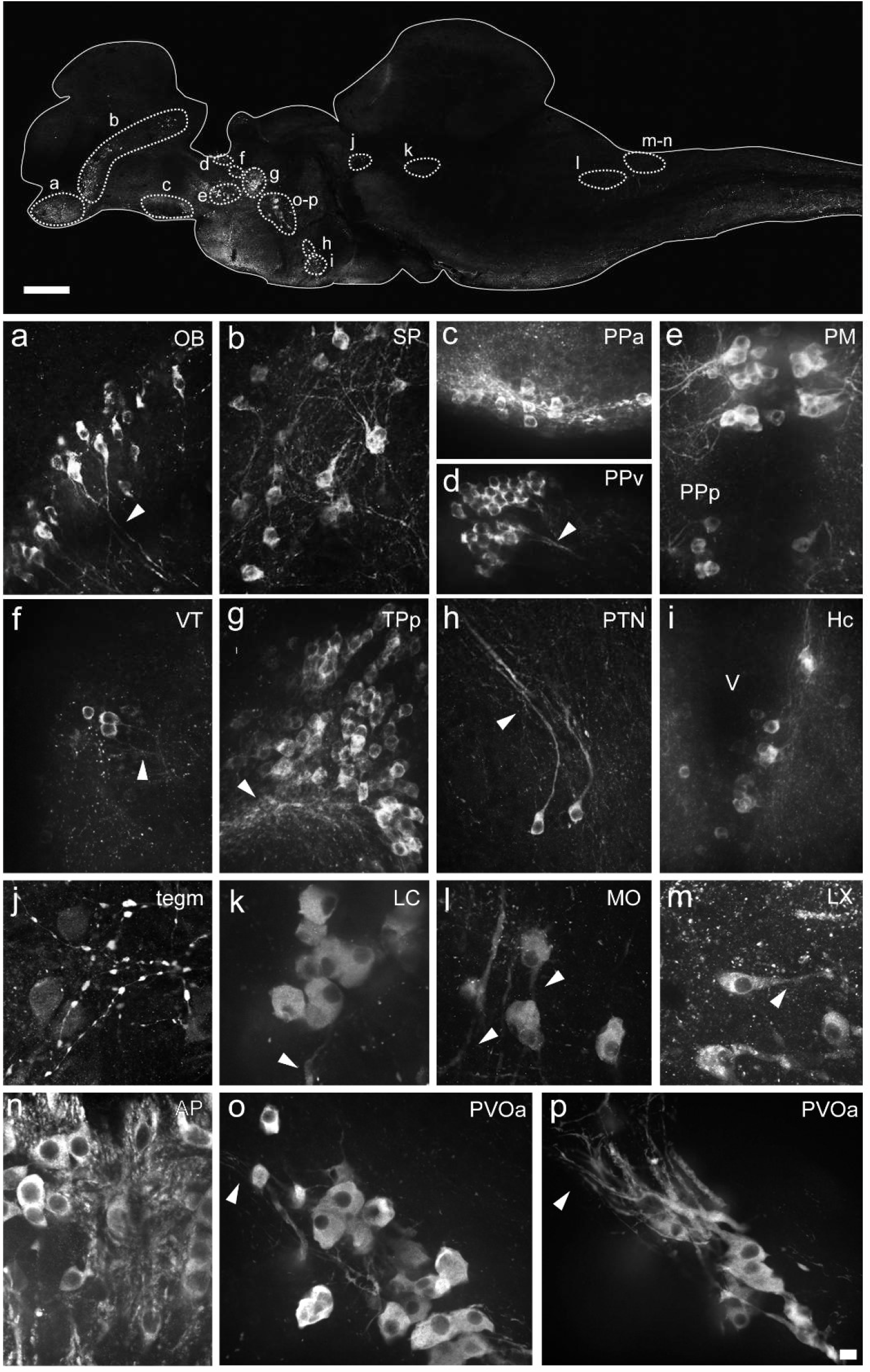
Morphology of the different types of TH+ neurons in the brain of *Nothobranchius furzeri*. The upper image illustrates the positions of TH+ cells in a para-sagittal section of the brain. (a-p) High resolution images of TH+ neurons present in the olfactory bulb (OB) **(A)**, subpallium (SP) **(B)**, anterior preoptic nucleus (PPa) **(C)**, ventral periventricular pretectal nucleus (PPv) **(D)**, magnocellular preoptic nucleus (PM) and parvocellular portion of preoptic nucleus (PPp) **(E)**, ventral thalamus (VT) **(F)**, periventricular nucleus of posterior tuberculum (TPp) **(G)**, posterior tuberal nucleus (PTN) **(H)**, caudal hypothalamus (Hc) **(I)**, mesencephalic tegmentum (Tegm) **(J)**, locus coeruleus (LC) **(K)**, medulla oblongata (IRF) **(L)**, vagal lobe (LX) **(M)**, area postrema (AP) **(N)**, and in the rostral **(O)** and caudal **(P)** clusters of the paraventricular organ-accompanying cells (PVOa). Arrowheads points to TH+ neuronal processes. For abbreviations see list. Scale bar, 500 µm (upper panel) and 10 µm (a-p).

**Figure 8.**
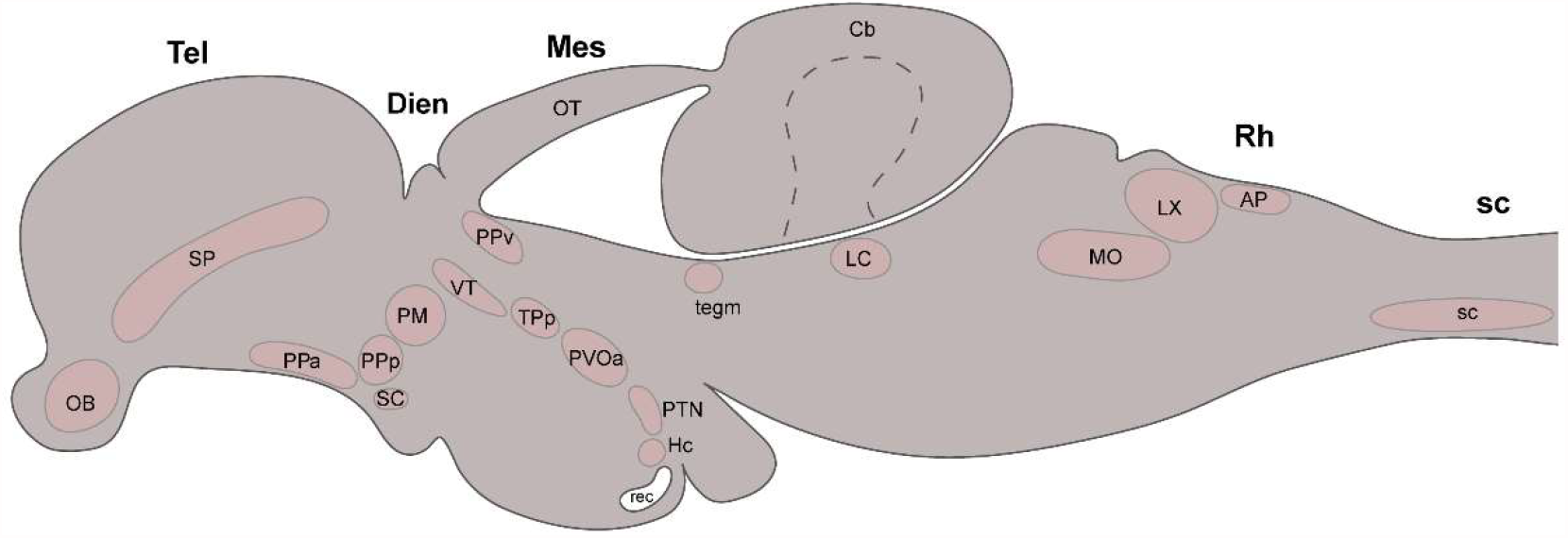
Summary diagram illustrating the localization of the catecholaminergic neuronal groups in a lateral view of the brain of *Nothobranchius furzeri*. For abbreviations see list.

## 4 Discussion

The distribution of catecholamines appears to be highly conserved throughout evolution and constitutes one of the oldest neurochemical systems in the vertebrate brain (Carlberg and Anctil, 1993; Yamamoto et al., 2011). Despite the variation in the morphology and complexity of the vertebrate brain, five main groups of CAergic cells can be identified in the CNS of all vertebrates. These conserved groups include the noradrenergic groups of the caudal and rostral/isthmic rhombencephalon and the dopaminergic groups located in the diencephalon, olfactory bulb and the retina, which suggests that certain essential functions have been preserved during evolution (Smeets and Gonzalez, 2000). However, other groups such as those located in the midbrain are more variable and while they are commonly found in the tetrapod lineage, they are unusual among actinopterygians. In contrast, the existence of DAergic groups in the subpallium and pretectum is abundant among actinopterygians but it is not a common feature of amniotes. Some of these inter-group differences has been attributed to brain function adaptations to the different types of vertebrate lifes (Yamamoto and Vernier, 2011).

Here we show that the CAergic system of *Nothobranchius furzeri* shares the general spatial organization observed among actinopterygians, in particular in the teleost group. However, the killifish shows some distinct features compared with other teleosts. Next, we discuss from a comparative perspective, the main conserved and divergent features of the organization of the CAergic system of *Nothobranchius furzeri*.

### 4.1 Telencephalon

#### Olfactory bulb

The rostralmost TH+ cells in the forebrain of *Nothobranchius furzeri* are found in the OB. The presence of DAergic cells in the OB is a constant feature among vertebrates and in mammals, according to the classical nomenclature, they correspond to the group A16 (Björklund and Lindvall, 1984; Hökfelt et al., 1984). In mice, dopamine exerts a negative regulatory role on olfactory inputs, inhibiting the release of glutamate to favor discrimination between odorants (Kruzich and Grandy, 2004). This function seems to be conserved in different taxa and has been described in fish such as goldfish (Kawai et al., 2012). The density and distribution of CAergic neurons in the layers of the OB is variable among vertebrates. While in mammals DAergic cells are more abundant in the glomerular and external layers, in teleosts they predominate in the deeper layers (Alonso et al., 1989; Batten et al., 1993; Smeets and Gonzalez, 2000). The latter organization is considered a primitive condition in vertebrates and is the most widespread among teleosts. However, there are several exceptions where cells adopt more external locations, such as in the case of zebrafish (Byrd and Brunjes, 1995), the ovoviviparous fishes *Poencilla reticulata* and *Gambusia affinis* (Bhat and Ganesh, 2017; Parafati et al., 2009), and the electric fish *Apteronotus leptorhynchus* (Sas et al., 1990). The origin of these cytoarchitectonic variations is unknown, but it appears to be a derived condition present in a few teleosts and possibly related to the way CAergic neurons integrate the chemosensitive information in these species (Parafati et al., 2009). In our study, we found that killifish TH+ cells adopt a peripheral position in the OB, being located primarily in the outermost layers. The significance of this distribution in the OB and whether it responds to an adaptation to the particular ecological niche of *Nothobranchius furzeri* remain to be investigated.

#### Telencephalic hemispheres

Similar to other teleosts, the killifish brain does not contain TH+ neurons in the pallium but it does in the subpallium. The presence of DAergic cells in the subpallium is an ancestral feature and has been described in cyclostomes, chondrichthyans and all actinopterygians (Yamamoto and Vernier, 2011). In contrast, these cells are absent in sarcopterygians such as lungfishes, and among tetrapods they have been only reported in monotremes and primates (Betarbet et al., 1997; Lopez and Gonzalez, 2017; Manger et al., 2002). The function of the subpallial CAergic neurons is unknown, but it has been proposed they project locally and provide DA to this region (Huot et al., 2007). Likewise, in the *Nothobranchius furzeri* a substantial fraction of the subpallial TH+ neurons showed confined projections to the ventral telencephalic area.

### 4.2 Diencephalon

#### Preoptic region

The distribution of CAergic neurons in the preoptic region of *Nothobranchius furzeri* is similar to other fish, including cyclostomes, chondrichthyans, actinopterygians and sarcopterygians (Bhat and Ganesh, 2017; Lopez and Gonzalez, 2017; Ma, 2003; Meredith and Smeets, 1987; Reiner and Northcutt, 1992; Wicht and Northcutt, 1994). In teleosts such as zebrafish, PO DAergic neurons send extensive projections to the pituitary resembling the tuberohypophyseal system of mammals (A14, A15) (Filippi et al., 2010; Kaslin and Panula, 2001). At a functional level, the most anteroventral TH+ cells of the PO (PPa) regulate the release of luteinizing hormone from the adenohypophysis (Bryant et al., 2016; Fontaine et al., 2015; Kah et al., 1984). Likewise, the PPa TH+ neurons of killifish send long-range projections to the ventral hypothalamus and apparently reach the hypophysis (**Figure S2B-C**). However, it remains to be determined whether these CAergic neurons also have a regulatory role in reproduction.

#### Ventral thalamus and pretectum

The thalamus of *Nothobranchius furzeri* contains TH+ neurons in its ventral subdivision (VT). DAergic cells are commonly found in this brain region in elasmobranchs, chondrosteans, cladistians, holosteans and teleosts (Adrio et al., 2002; Lopez et al., 2019; Lozano et al., 2019). In mammals, a comparable population of CAergic cells are located in the zona incerta and corresponds to the A13 group (Meister et al., 1987; Smeets and Gonzalez, 2000). Remarkably, DAergic neurons in the VT of zebrafish and the prethalamic groups described in mammals share the same transcription factors (Arx and Isl1) during development suggesting they are homologous groups. However, at the functional level such homology is less clear (Filippi et al., 2012).

In the pretectal region (PPv) of *Nothobranchius furzeri* exists conspicuous group of CAergic neurons with dendritic extension to the optic tectum. The pretectal DAergic cells are frequently found in bony fishes and amniotes, with the exception of mammals (Smeets and Gonzalez, 2000). These cells could be implicated in the modulation of the retino-tectal visual input (Yamamoto and Vernier, 2011).

#### Posterior tuberculum and hypothalamus

*Nothobranchius furzeri* exhibits different groups of CAergic neurons in the posterior tuberculum that are similar in terms of morphology and projection pattern to other teleosts (Bhat and Ganesh, 2017; Forlano and Sisneros, 2016; Karoubi et al., 2016; Kaslin and Panula, 2001; Rink and Wullimann, 2001) and other groups of fish such as cyclostomes (Pierre et al., 1997), elasmobranchs (Molist et al., 1993), holosteans (Lozano et al., 2019), chondrosteans (Adrio et al., 2002) and cladistians (Lopez et al., 2019; Reiner and Northcutt, 1992). Studies in zebrafish and in *Polypterus senegalus* demonstrated that some of these neurons project onto a region homologous to the mammalian striatum (Reiner and Northcutt, 1992; Rink and Wullimann, 2001; Tay et al., 2011) and degenerate in toxic-induced models of Parkinson’s Disease (Sallinen et al., 2009). Furthermore, zebrafish TP neurons require NR4A2/Nurr1 for their development, in the same way that DAergic neurons of the mammalian substantia nigra/ventral tegmental area (SN/VTA) do (Blin et al., 2008). These findings, added to the lack of DAergic groups in the mesencephalon of actinopterygians, have led to the proposal that part of the neurons of the TP correspond to the A9/A10 groups of mammals. On the other hand, a subgroup of large TH+ cells in the posterior tuberculum of zebrafish projects to the hindbrain and spinal cord, and modulate motor and sensory activity (Reinig et al., 2017; Tay et al., 2011). In addition, these neurons require Nkx2.1 and Otp for their differentiation as also observed for the A11 neurons of mice (Ryu et al., 2007). Thereby, it has been suggested that different subpopulations of DAergic neurons that coexist in the posterior tuberculum of teleosts resemble the A9/A10 and A11 groups of mammals (Yamamoto and Vernier, 2011). Consistent with this proposal, we found that killifish contains different subgroups of TH+ neurons in the posterior tuberculum that either send ascending projections in direction to the telencephalon or descending projections to the medulla oblongata. Previously, Matsui et al. (2019) reported that the large TH+ neurons located in the TP suffer an age-dependent degeneration and accumulate α-synuclein aggregates, thus reinforcing the idea that this group shares similar properties to the DAergic neurons of the mammalian substantia nigra.

The distribution of TH+ cells in the hypothalamus of *Nothobranchius furzeri* seems to be less conserved compared to other teleosts. The killifish lacks TH immunoreactivity in the PVO and only has positive neurons in the caudal hypothalamus (Hc). The presence of cerebrospinal fluid-contacting cells (CSF-c) containing monoamine in the PVO occurs in most vertebrates, but is reduced in the tetrapod lineage and even absent in placental mammals (Xavier et al., 2017). In zebrafish, the PVO CSF-c mainly express the *th2* transcript of the tyrosine hydroxylase enzyme, however, a group of neurons located in the most rostral region of the PVO expresses *th1* (Yamamoto et al., 2010). By contrast, the holosteans and sarcopterygians also contain DAergic cells in the PVO, but they use the *th2* enzyme and thus lack immunoreactivity with common TH antibodies (Lopez and Gonzalez, 2017; Lozano et al., 2019). We found that the PVO of killifish is TH negative, but we cannot rule out the existence of *th2* DAergic CSF-c in this region. The functional significance of DAergic neurons in the PVO is not clear, but they seem to play a role in deep brain photoreception in non-mammalian vertebrates (Vigh et al., 2002). Furthermore, unlike other teleosts that present a more extended TH labeling in the basal hypothalamus (Rink and Wullimann, 2001), the CAergic neurons of killifish are restricted to the Hc. Likewise as the PVO, the TH+ cells in the Hc are absent in mammals and only are observed in some groups of actinopterygians (Xavier et al., 2017). In zebrafish, two different population of CSF-c co-exist in this region, one that expresses *th1* and the other *th2* gene. The latter group seems to play a specific role in fishes associated with the initiation of swimming behavior (McPherson et al., 2016). Instead, the *th1* cluster appears to have a functional correlate with the A14 group of mammals, based on their projections (Tay et al., 2011). The TH+ cells in the Hc of killifish also send local processes on the ventral hypothalamus as seen for the Hc cluster in zebrafish. Thus, it is possible that this group also corresponds to the A14 cells of the mammals.

### 4.3 Mesencephalon

Strikingly, a small group of lightly stained TH+ cells is detected in the rostral mesencephalic tegmentum of *Nothobranchius furzeri*, near the ventricle and dorsal to the oculomotor nucleus. The absence of DAergic neurons in the mesencephalic tegmentum is a common trait of cyclostomes and most species of actinopterygians (Lopez et al., 2019), with a few exceptions that include the holostean *Amia calva* and *Lepisosteus osseus* (Lozano et al., 2019; Parent and Northcutt, 1982), the cladistia *Polypterus senegalus* (Lopez et al., 2019) and the teleost *Anguilla Anguilla* (Roberts et al., 1989). This mesencephalic group differs in terms of location and morphology from the SN/VTA found in the elasmobranchs, lungfish and tetrapods (Lopez and Gonzalez, 2017; Smeets and Gonzalez, 2000). The connectivity and functional significance of this CAergic cluster is still unknown.

### 4.4 Rhombencephalon

#### Dorsal rhombencephalon

In the rostral rhombencephalon, *Nothobranchius furzeri* shows a small population of large TH+ cells showing ventral projections that correspond to the locus coeruleus (LC). These cells constitute a conserved CAergic groups among vertebrates and is one of the main sources of noradrenaline (NA) in the CNS (Smeets and Gonzalez, 2000). In mammals, the LC is recognized as the A6 group and innervates widely the brain and the spinal cord. Given the complex and vast projections of the LC, this nucleus is associated with a wide range of functions including cardiovascular control, olfactory and auditory information processing, endocrine control, nociception, arousal state and motor behavior, among others (Aston-Jones and Bloom, 1981). A similar connectivity has been described for the LC in teleosts (Ma, 1994a, b; Tay et al., 2011). Although the role of this noradrenergic (NAergic) group is less clear in fishes, a recent study in zebrafish shows that the LC mediates wakefulness through the hypocretin-induced arousal system (Singh et al., 2015).

#### Medulla oblongata

Three groups of TH+ neurons are observed in the caudal rhombencephalon of *Nothobranchius furzeri*, along the medulla oblongata spanning the vagal lobe and the area postrema. Comparable NAergic neurons have been described in the caudal hindbrain of other teleosts, holosteans, cladistians and lungfish (Kaslin and Panula, 2001; Lopez and Gonzalez, 2017; Lopez et al., 2019; Lozano et al., 2019; Ma, 1997). In mammals, the NAergic neurons of the medulla oblongata correspond to the A1-A2 group and are involved in sympathetic functions such as the control of arterial pressure, respiratory peacemaking and response to pH (Freiria-Oliveira et al., 2015). It has been proposed that the NAergic neurons in the caudal rombencephalon have a conserved function in vertebrates (Smeets and Gonzalez, 2000).

### 4.5 Spinal cord

The ventral aspect of the spinal cord (sc) in *Nothobranchius furzeri* houses small TH+ neurons, located away from the central canal. Two types of sc DAergic cells have been described in vertebrates, one with CSF-c characteristics and the other lacking contacts with the central canal. The former type is widely distributed in fish and has been reported in cyclostomes (lampreys), elasmobranchs (rays), teleosts (not in zebrafish), holosteans, cladistians and lungfish, but are absent in mammals (Lopez and Gonzalez, 2017; Lopez et al., 2019; Lozano et al., 2019; Roberts and Meredith, 1987; Rodicio et al., 2008; Sueiro et al., 2004). The second type has only been detected in lampreys, the eel *Anguilla anguilla* and in a few mammals as rats and monkeys (Roberts et al., 1989; Rodicio et al., 2008; Smeets and Gonzalez, 2000). The significance of CAergic neurons in the spinal cord is not fully understood, but could contribute to modulation of spinal reflexes, nociception and locomotion (Smeets and Gonzalez, 2000). The TH+ sc neurons in the killifish do not appear to contact the central canal and it remains to be determined whether they correspond to the second neuronal type described in other species.

### 4.6 Retina

The retina of *Nothobranchius furzeri* displays a discrete number of TH+ neurons in the INL, which according to their morphology appear to correspond to amacrine cells (Gatta et al., 2014). The presence of TH+ amacrine cells has been described in almost all vertebrates studied to date and in mammals constitute the group A17. The DAergic amacrine cells play a conserved role in retinal adaptation to light through the regulation of retinomotor movements and changes in the size of the receptive field (Masland, 2001).

### 4.7 Concluding remarks

*Nothobranchius furzeri* belongs to the teleosts, one of the largest taxa among vertebrates with large variability in brain neuroanatomical structure as a result of extreme specialized behaviors. Despite this variability, teleost fish have a common organization of the CArgic system that in turn is largely conserved with other vertebrates. In this context, killifish shares the same basic pattern seen in other teleosts. However, unlike other members of this group such as zebrafish, killifish shows a more limited number of TH+ groups in the hypothalamus and has CAergic neurons in the spinal cord. Furthermore, the killifish shows a particular group of CAergic neurons in the mesencephalic tegmentum that is an unusual trait among actinopterygians. Future studies will provide a more detailed analysis of the ontogeny, neurotransmitter phenotype (DA, NA) and the afferent/efferent connections of the different TH+ groups of *Nothobranchius furzeri*, to dissect the functional correlate with CAergic neuronal groups of other species.

## Ethics Statement

The animal study was reviewed and approved by the Bioethics Committee of the Faculty of Medicine, University of Chile (CICUA certificate number: 20385-MED-UCH).

## Author Contributions

JB and AOP performed the tissue processing and immunofluorescence assays. JB, PAG, AOP, CAC, PH, and MLC contributed to the analysis of the data and created the figures. MLC supervised the work. JB and MLC wrote the manuscript with input from PAG and PH.

## Abbreviations

AP: area postrema
Cant: commissura anterior
Cb: cerebellum
CC: cerebellar crest
CCe: corpus of cerebellum
Dc1-2-3-4: central zone of dorsal telencephalon 1-4
Dd: dorsal zone of dorsal telencephalon
Dien: diencephalon
DIL: diffuse nucleus of inferior lobe
Dld: dorso-lateral zone of dorsal telencephalon
Dll: laterol-lateral zone of dorsal telencephalon
Dlv: ventro-lateral zone of dorsal telencephalon
Dm1-2-3-4: medial zone of dorsal telencephalon 1-4
ECL: external cellular layer
Ed: dorsal entopeduncular nucleus
EG: granular eminence
gc: griseum centrale
GCL: ganglion cell layer
gl: glomerular layer
Gl: glomerular layer of the olfactory bulb
Ha: habenular nucleus
Hc: caudal hypothalamus
Hv: ventral hypothalamus
ICL: internal cellular layer
INL: inner nuclear layer
IPL: inner plexiform layer
LC: locus coeruleus
lt: longitudinal tract
lfb: lateral forebrain bundle
llf: lateral longitudinal fascicle
IMRF: intermediate reticular formation
IRF: inferior reticular formation
LVII: facial lobe
LX: vagal lobe
Mes: mesencephalon
mlf: medial longitudinal fascicle
MO: medulla oblongata
MON: medial octavolateralis nucleus
NG: glomerular nucleus
NIII: oculomotor nucleus
nV: nerve V
OB: olfactory bulb
ON: optic nerve
ONL: outer nuclear layer
OPL: outer plexiform layer
OS: outer segment
OT: optic tectum
PGc: central preglomerular nucleus
PGZ: periglomerular gray zone
PM: magnocellular preoptic nucleus
PO: preoptic area
PTN: posterior tuberal nucleus
poht: preopticohypothalamic tract
PPa: anterior preoptic nucleus
PPp: parvocellular portion of preoptic nucleus
PPv: ventral periventricular pretectal nucleus
PVO: paraventricular organ
PVOa: paraventricular organ-accompanying cells
rec: posterior recess
Rh: rhomboenchephalon
sc: spinal cord
SC: suprachiasmatic nucleus
SP: subpallium
SRF: superior reticular formation
tegm: mesencephalic tegmentum
Tel: telencephalon
Tl: torus longitudinalis
TPp: periventricular nucleus of posterior tuberculum
TS: torus semicircularis
V: ventricle
Va: valvula of cerebellum
Vd: dorsal zone of ventral telencephalon
Vp: posterior zone of ventral telencephalon
Vs: supracommissural zone of ventral telencephalon
VT: ventral thalamic nucleus
Vv: ventral zone of ventral telencephalon

## Acknowledgments

We thank the support of the annual killifish facility of the Faculty of Medicine, University of Chile, jointly funded by ICBM, GERO and BNI. This work was funded by the following projects of the Chilean National Agency for Research and Development (ANID): Millennium Institute ICN09_015 to M.L.C and P.A-G.; FONDAP 15150012 to M.L.C., J.B and C.A-C.; FONDEQUIP EQM130051 to M.L.C; and Ring PIA ACT-192015 to M.L.C.

## References

Adrio, F., Anadon, R., Rodriguez-Moldes, I. (2002). Distribution of tyrosine hydroxylase (TH) and dopamine beta-hydroxylase (DBH) immunoreactivity in the central nervous system of two chondrostean fishes (Acipenser baeri and Huso huso). The Journal of comparative neurology 448, 280–297.

Alonso, J.R., Covenas, R., Lara, J., Arevalo, R., de Leon, M., Aijon, J., 1989. Tyrosine hydroxylase immunoreactivity in a subpopulation of granule cells in the olfactory bulb of teleost fish. Brain Behav Evol 34, 318–324.

Aston-Jones, G., Bloom, F.E., 1981. Activity of norepinephrine-containing locus coeruleus neurons in behaving rats anticipates fluctuations in the sleep-waking cycle. The Journal of neuroscience : the official journal of the Society for Neuroscience 1, 876–886.

Batten, T.F., Berry, P.A., Maqbool, A., Moons, L., Vandesande, F., 1993. Immunolocalization of catecholamine enzymes, serotonin, dopamine and L-dopa in the brain of Dicentrarchus labrax (Teleostei). Brain Res Bull 31, 233–252.

Baumgart, M., Di Cicco, E., Rossi, G., Cellerino, A., Tozzini, E.T., 2015. Comparison of captive lifespan, age-associated liver neoplasias and age-dependent gene expression between two annual fish species: Nothobranchius furzeri and Nothobranchius korthause. Biogerontology 16, 63–69.

Benavides-Piccione, R., DeFelipe, J., 2007. Distribution of neurons expressing tyrosine hydroxylase in the human cerebral cortex. J Anat 211, 212–222.

Bertler, A., Rosengren, E., 1959. Occurrence and distribution of catechol amines in brain. Acta Physiol Scand 47, 350–361.

Betarbet, R., Turner, R., Chockkan, V., DeLong, M.R., Allers, K.A., Walters, J., Levey, A.I., Greenamyre, J.T., 1997. Dopaminergic neurons intrinsic to the primate striatum. The Journal of neuroscience : the official journal of the Society for Neuroscience 17, 6761–6768.

Bhat, S.K., Ganesh, C.B., 2017. Distribution of tyrosine hydroxylase-immunoreactive neurons in the brain of the viviparous fish Gambusia affinis. J Chem Neuroanat 85, 1–12.

Björklund, A., Lindvall, O., 1984. Dopamine-containing system in the CNS, in: Björklund, A., Hökfelt, T. (Eds.), Handbook of Chemical Neuroanatomy Elsevier, Amsterdam, pp. 55–102.

Blazek, R., Polacik, M., Reichard, M., 2013. Rapid growth, early maturation and short generation time in African annual fishes. Evodevo 4, 24.

Blin, M., Norton, W., Bally-Cuif, L., Vernier, P., 2008. NR4A2 controls the differentiation of selective dopaminergic nuclei in the zebrafish brain. Mol Cell Neurosci 39, 592–604.

Brinon, J.G., Arevalo, R., Weruaga, E., Crespo, C., Alonso, J.R., Aijon, J., 1998. Tyrosine hydroxylase-like immunoreactivity in the brain of the teleost fish Tinca tinca. Arch Ital Biol 136, 17–44.

Bryant, A.S., Greenwood, A.K., Juntti, S.A., Byrne, A.E., Fernald, R.D., 2016. Dopaminergic inhibition of gonadotropin-releasing hormone neurons in the cichlid fish Astatotilapia burtoni. J Exp Biol 219, 3861–3865.

Byrd, C.A., Brunjes, P.C., 1995. Organization of the olfactory system in the adult zebrafish: histological, immunohistochemical, and quantitative analysis. The Journal of comparative neurology 358, 247–259.

Carlsson, A., 1959. The occurrence, distribution and physiological role of catecholamines in the nervous system. Pharmacol Rev 11, 490–493.

Carlberg, M., Anctil, M., 1993. Biogenic amines in coelenterates. Comp. Biochem. Physiol. C Comp. Pharmacol. Toxicol. 106, 1–9.

Crocker, A.D., 1997. The regulation of motor control: an evaluation of the role of dopamine receptors in the substantia nigra. Reviews in the neurosciences 8, 55–76.

D’Angelo, L., 2013. Brain atlas of an emerging teleostean model: Nothobranchius furzeri. Anat Rec (Hoboken) 296, 681–691.

Davis, M., Rainnie, D., Cassell, M., 1994. Neurotransmission in the rat amygdala related to fear and anxiety. Trends in neurosciences 17, 208–214.

Dawson, T.M., Dawson, V.L., 2003. Molecular pathways of neurodegeneration in Parkinson’s disease. Science 302, 819–822.

Di Cicco, E., Tozzini, E.T., Rossi, G., Cellerino, A., 2011. The short-lived annual fish Nothobranchius furzeri shows a typical teleost aging process reinforced by high incidence of age-dependent neoplasias. Exp Gerontol 46, 249–256.

Dubach, M., Schmidt, R., Kunkel, D., Bowden, D.M., Martin, R., German, D.C., 1987. Primate neostriatal neurons containing tyrosine hydroxylase: immunohistochemical evidence. Neurosci Lett 75, 205–210.

El Massri, N., Lemgruber, A.P., Rowe, I.J., Moro, C., Torres, N., Reinhart, F., Chabrol, C., Benabid, A.L., Mitrofanis, J., 2017. Photobiomodulation-induced changes in a monkey model of Parkinson’s disease: changes in tyrosine hydroxylase cells and GDNF expression in the striatum. Experimental brain research 235, 1861–1874.

Filippi, A., Jainok, C., Driever, W., 2012. Analysis of transcriptional codes for zebrafish dopaminergic neurons reveals essential functions of Arx and Isl1 in prethalamic dopaminergic neuron development. Dev Biol 369, 133–149.

Filippi, A., Mahler, J., Schweitzer, J., Driever, W., 2010. Expression of the paralogous tyrosine hydroxylase encoding genes th1 and th2 reveals the full complement of dopaminergic and noradrenergic neurons in zebrafish larval and juvenile brain. The Journal of comparative neurology 518, 423–438.

Fontaine, R., Affaticati, P., Bureau, C., Colin, I., Demarque, M., Dufour, S., Vernier, P., Yamamoto, K., Pasqualini, C., 2015. Dopaminergic Neurons Controlling Anterior Pituitary Functions: Anatomy and Ontogenesis in Zebrafish. Endocrinology 156, 2934–2948.

Forlano, P.M., Sisneros, J.A., 2016. Neuroanatomical Evidence for Catecholamines as Modulators of Audition and Acoustic Behavior in a Vocal Teleost. Adv Exp Med Biol 877, 439–475.

Freiria-Oliveira, A.H., Blanch, G.T., Pedrino, G.R., Cravo, S.L., Murphy, D., Menani, J.V., Colombari, D.S., 2015. Catecholaminergic neurons in the comissural region of the nucleus of the solitary tract modulate hyperosmolality-induced responses. Am J Physiol Regul Integr Comp Physiol 309, R1082–1091.

Gallman, K., Rivera, D., Soares, D., 2019. Evolutionary increases in catecholamine signaling may underlie the emergence of adaptive traits and behaviors in the blind cavefish, <em>Astyanax mexicanus</em>. bioRxiv, 724179.

Gatta, C., Castaldo, L., Cellerino, A., de Girolamo, P., Lucini, C., D’Angelo, L., 2014. Brain derived neurotrophic factor in the retina of the teleost N. furzeri. Ann Anat 196, 192–196.

Genade, T., Benedetti, M., Terzibasi, E., Roncaglia, P., Valenzano, D.R., Cattaneo, A., Cellerino, A., 2005. Annual fishes of the genus Nothobranchius as a model system for aging research. Aging Cell 4, 223–233.

Goebrecht, G.K., Kowtoniuk, R.A., Kelly, B.G., Kittelberger, J.M., 2014. Sexually-dimorphic expression of tyrosine hydroxylase immunoreactivity in the brain of a vocal teleost fish (Porichthys notatus). J Chem Neuroanat 56, 13–34.

Hökfelt, T., Martesson, R., Björklund, A., Kleinau, S., Goldstein, M., 1984. Distributional maps of tyrosine-hydroxylase-immunoreactive neurons in the rat brain., in: Björklund, A., Hökfelt, T. (Eds.), Classical Transmitters in the CNS, Part I, Handbook of Chemical Neuroanatomy Elsevier, Amsterdam, pp. 277–386.

Hornby, P.J., Piekut, D.T., 1990. Distribution of catecholamine-synthesizing enzymes in goldfish brains: presumptive dopamine and norepinephrine neuronal organization. Brain Behav Evol 35, 49–64.

Huisman, E., Uylings, H.B., Hoogland, P.V., 2004. A 100% increase of dopaminergic cells in the olfactory bulb may explain hyposmia in Parkinson’s disease. Movement disorders : official journal of the Movement Disorder Society 19, 687–692.

Huisman, E., Uylings, H.B., Hoogland, P.V., 2008. Gender-related changes in increase of dopaminergic neurons in the olfactory bulb of Parkinson’s disease patients. Movement disorders : official journal of the Movement Disorder Society 23, 1407–1413.

Huot, P., Levesque, M., Parent, A., 2007. The fate of striatal dopaminergic neurons in Parkinson’s disease and Huntington’s chorea. Brain : a journal of neurology 130, 222–232.

Hurley, L.M., Devilbiss, D.M., Waterhouse, B.D., 2004. A matter of focus: monoaminergic modulation of stimulus coding in mammalian sensory networks. Current opinion in neurobiology 14, 488–495.

Joyce, J.N., 1993. The dopamine hypothesis of schizophrenia: limbic interactions with serotonin and norepinephrine. Psychopharmacology 112, S16–34.

Kah, O., Chambolle, P., Thibault, J., Geffard, M., 1984. Existence of dopaminergic neurons in the preoptic region of the goldfish. Neurosci Lett 48, 293–298.

Karoubi, N., Segev, R., Wullimann, M.F., 2016. The Brain of the Archerfish Toxotes chatareus: A Nissl-Based Neuroanatomical Atlas and Catecholaminergic/Cholinergic Systems. Front Neuroanat 10, 106.

Kaslin, J., Panula, P., 2001. Comparative anatomy of the histaminergic and other aminergic systems in zebrafish (Danio rerio). The Journal of comparative neurology 440, 342–377.

Kawai, T., Abe, H., Oka, Y., 2012. Dopaminergic neuromodulation of synaptic transmission between mitral and granule cells in the teleost olfactory bulb. J Neurophysiol 107, 1313–1324.

Kruzich, P.J., Grandy, D.K., 2004. Dopamine D2 receptors mediate two-odor discrimination and reversal learning in C57BL/6 mice. BMC Neurosci 5, 12.

Lopez, J.M., Gonzalez, A., 2017. Organization of the catecholaminergic systems in the brain of lungfishes, the closest living relatives of terrestrial vertebrates. The Journal of comparative neurology 525, 3083–3109.

Lopez, J.M., Lozano, D., Morona, R., Gonzalez, A., 2019. Organization of the catecholaminergic systems in two basal actinopterygian fishes, Polypterus senegalus and Erpetoichthys calabaricus (Actinopterygii: Cladistia). The Journal of comparative neurology 527, 437–461.

Lozano, D., Morona, R., Gonzalez, A., Lopez, J.M., 2019. Comparative Analysis of the Organization of the Catecholaminergic Systems in the Brain of Holostean Fishes (Actinopterygii/Neopterygii). Brain Behav Evol 93, 206–235.

Ma, P.M., 1994a. Catecholaminergic systems in the zebrafish. I. Number, morphology, and histochemical characteristics of neurons in the locus coeruleus. The Journal of comparative neurology 344, 242–255.

Ma, P.M., 1994b. Catecholaminergic systems in the zebrafish. II. Projection pathways and pattern of termination of the locus coeruleus. The Journal of comparative neurology 344, 256–269.

Ma, P.M., 1997. Catecholaminergic systems in the zebrafish. III. Organization and projection pattern of medullary dopaminergic and noradrenergic neurons. The Journal of comparative neurology 381, 411–427.

Ma, P.M., 2003. Catecholaminergic systems in the zebrafish. IV. Organization and projection pattern of dopaminergic neurons in the diencephalon. The Journal of comparative neurology 460, 13–37.

Manaye, K.F., McIntire, D.D., Mann, D.M., German, D.C., 1995. Locus coeruleus cell loss in the aging human brain: a non-random process. The Journal of comparative neurology 358, 79–87.

Manger, P.R., Fahringer, H.M., Pettigrew, J.D., Siegel, J.M., 2002. The distribution and morphological characteristics of catecholaminergic cells in the brain of monotremes as revealed by tyrosine hydroxylase immunohistochemistry. Brain Behav Evol 60, 298–314.

Manso, M.J., Becerra, M., Molist, P., Rodriguez-Moldes, I., Anadon, R., 1993. Distribution and development of catecholaminergic neurons in the brain of the brown trout. A tyrosine hydroxylase immunohistochemical study. J Hirnforsch 34, 239–260.

Masland, R.H., 2001. The fundamental plan of the retina. Nat Neurosci 4, 877–886.

Matsui, H., Kenmochi, N., Namikawa, K., 2019. Age-and alpha-Synuclein-Dependent Degeneration of Dopamine and Noradrenaline Neurons in the Annual Killifish Nothobranchius furzeri. Cell Rep 26, 1727–1733 e1726.

Matsui, H., Sugie, A., 2017. An optimized method for counting dopaminergic neurons in zebrafish. PloS one 12, e0184363.

Matsui, H., Takahashi, R., 2018. Parkinson’s disease pathogenesis from the viewpoint of small fish models. Journal of neural transmission 125, 25–33.

Matsui, H., Uemura, N., Yamakado, H., Takeda, S., Takahashi, R., 2014. Exploring the pathogenetic mechanisms underlying Parkinson’s disease in medaka fish. J Parkinsons Dis 4, 301–310.

McPherson, A.D., Barrios, J.P., Luks-Morgan, S.J., Manfredi, J.P., Bonkowsky, J.L., Douglass, A.D., Dorsky, R.I., 2016. Motor Behavior Mediated by Continuously Generated Dopaminergic Neurons in the Zebrafish Hypothalamus Recovers after Cell Ablation. Curr Biol 26, 263–269.

Meek, J., Joosten, H.W., 1993. Tyrosine hydroxylase-immunoreactive cell groups in the brain of the teleost fish Gnathonemus petersii. J Chem Neuroanat 6, 431–446.

Meister, B., Hokfelt, T., Brown, J., Joh, T., Goldstein, M., 1987. Dopaminergic cells in the caudal A13 cell group express somatostatin-like immunoreactivity. Experimental brain research 67, 441–444.

Menuet, A., Alunni, A., Joly, J.S., Jeffery, W.R., Retaux, S., 2007. Expanded expression of Sonic Hedgehog in Astyanax cavefish: multiple consequences on forebrain development and evolution. Development 134, 845–855.

Meredith, G.E., Smeets, W.J., 1987. Immunocytochemical analysis of the dopamine system in the forebrain and midbrain of Raja radiata: evidence for a substantia nigra and ventral tegmental area in cartilaginous fish. The Journal of comparative neurology 265, 530–548.

Molist, P., Rodriguez-Moldes, I., Anadon, R., 1993. Organization of catecholaminergic systems in the hypothalamus of two elasmobranch species, Raja undulata and Scyliorhinus canicula. A histofluorescence and immunohistochemical study. Brain Behav Evol 41, 290–302.

Montagu, K.A., 1956. Adrenaline and noradrenaline concentrations in rat tissues. Biochem J 63, 559–565.

Northcutt, R.G., Reiner, A., Karten, H.J., 1988. Immunohistochemical study of the telencephalon of the spiny dogfish, Squalus acanthias. The Journal of comparative neurology 277, 250–267.

Palfi, S., Leventhal, L., Chu, Y., Ma, S.Y., Emborg, M., Bakay, R., Deglon, N., Hantraye, P., Aebischer, P., Kordower, J.H., 2002. Lentivirally delivered glial cell line-derived neurotrophic factor increases the number of striatal dopaminergic neurons in primate models of nigrostriatal degeneration. The Journal of neuroscience : the official journal of the Society for Neuroscience 22, 4942–4954.

Parafati, M., Senatori, O., Nicotra, A., 2009. Localization of tyrosine hydroxylase immunoreactive neurons in the forebrain of the guppy Poecilia reticulata. J Fish Biol 75, 1194–1205.

Parent, A., Northcutt, R.G., 1982. The monoamine-containing neurons in the brain of the garfish, Lepisosteus osseus. Brain Res Bull 9, 189–204.

Pierre, J., Mahouche, M., Suderevskaya, E.I., Reperant, J., Ward, R., 1997. Immunocytochemical localization of dopamine and its synthetic enzymes in the central nervous system of the lamprey Lampetra fluviatilis. The Journal of comparative neurology 380, 119–135.

Puelles, L. & Rubenstein, J. L., 2003. Forebrain gene expression domains and the evolving prosomeric model. Trends Neurosciences 26, 469–476.

Reichard, M., Polacik, M., Sedlacek, O., 2009. Distribution, colour polymorphism and habitat use of the African killifish Nothobranchius furzeri, the vertebrate with the shortest life span. J Fish Biol 74, 198–212.

Reiner, A., Northcutt, R.G., 1992. An immunohistochemical study of the telencephalon of the senegal bichir (Polypterus senegalus). The Journal of comparative neurology 319, 359–386.

Reinig, S., Driever, W., Arrenberg, A.B., 2017. The Descending Diencephalic Dopamine System Is Tuned to Sensory Stimuli. Curr Biol 27, 318–333.

Rink, E., Wullimann, M.F., 2001. The teleostean (zebrafish) dopaminergic system ascending to the subpallium (striatum) is located in the basal diencephalon (posterior tuberculum). Brain research 889, 316–330.

Roberts, B.L., Meredith, G.E., 1987. Immunohistochemical study of a dopaminergic system in the spinal cord of the ray, Raja radiata. Brain research 437, 171–175.

Roberts, B.L., Meredith, G.E., Maslam, S., 1989. Immunocytochemical analysis of the dopamine system in the brain and spinal cord of the European eel, Anguilla anguilla. Anat Embryol (Berl) 180, 401–412.

Rodicio, M.C., Villar-Cervino, V., Barreiro-Iglesias, A., Anadon, R., 2008. Colocalization of dopamine and GABA in spinal cord neurones in the sea lamprey. Brain Res Bull 76, 45–49.

Rodriguez-Gomez, F.J., Rendon-Unceta, M.C., Sarasquete, C., Munoz-Cueto, J.A., 2000. Localization of tyrosine hydroxylase-immunoreactivity in the brain of the Senegalese sole, Solea senegalensis. J Chem Neuroanat 19, 17–32.

Roufail, E., Rees, S., 1997. Ageing has a differential effect on nitric oxide synthase-containing and catecholaminergic amacrine cells in the human and rat retina. The Journal of comparative neurology 389, 329–347.

Ryu, S., Mahler, J., Acampora, D., Holzschuh, J., Erhardt, S., Omodei, D., Simeone, A., Driever, W., 2007. Orthopedia homeodomain protein is essential for diencephalic dopaminergic neuron development. Curr Biol 17, 873–880.

Salamone, J.D., Correa, M., 2012. The mysterious motivational functions of mesolimbic dopamine. Neuron 76, 470–485.

Sallinen, V., Torkko, V., Sundvik, M., Reenila, I., Khrustalyov, D., Kaslin, J., Panula, P., 2009. MPTP and MPP+ target specific aminergic cell populations in larval zebrafish. J Neurochem 108, 719–731.

Sas, E., Maler, L., Tinner, B., 1990. Catecholaminergic systems in the brain of a gymnotiform teleost fish: an immunohistochemical study. The Journal of comparative neurology 292, 127–162.

Sheng, D., Qu, D., Kwok, K.H., Ng, S.S., Lim, A.Y., Aw, S.S., Lee, C.W., Sung, W.K., Tan, E.K., Lufkin, T., Jesuthasan, S., Sinnakaruppan, M., Liu, J., 2010. Deletion of the WD40 domain of LRRK2 in Zebrafish causes Parkinsonism-like loss of neurons and locomotive defect. PLoS Genet 6, e1000914.

Singh, C., Oikonomou, G., Prober, D.A., 2015. Norepinephrine is required to promote wakefulness and for hypocretin-induced arousal in zebrafish. Elife 4, e07000.

Smeets, W.J., Gonzalez, A., 2000. Catecholamine systems in the brain of vertebrates: new perspectives through a comparative approach. Brain Res Brain Res Rev 33, 308–379.

Soman, S.K., Bazala, M., Keatinge, M., Bandmann, O., Kuznicki, J., 2019. Restriction of mitochondrial calcium overload by mcu inactivation renders a neuroprotective effect in zebrafish models of Parkinson’s disease. Biol Open 8.

Sueiro, C., Carrera, I., Molist, P., Rodriguez-Moldes, I., Anadon, R., 2004. Distribution and development of glutamic acid decarboxylase immunoreactivity in the spinal cord of the dogfish Scyliorhinus canicula (elasmobranchs). The Journal of comparative neurology 478, 189–206.

Tande, D., Hoglinger, G., Debeir, T., Freundlieb, N., Hirsch, E.C., Francois, C., 2006. New striatal dopamine neurons in MPTP-treated macaques result from a phenotypic shift and not neurogenesis. Brain : a journal of neurology 129, 1194–1200.

Tashiro, Y., Kaneko, T., Nagatsu, I., Kikuchi, H., Mizuno, N., 1990. Increase of tyrosine hydroxylase-like immunoreactive neurons in the nucleus accumbens and the olfactory bulb in the rat with the lesion in the ventral tegmental area of the midbrain. Brain research 531, 159–166.

Tashiro, Y., Sugimoto, T., Hattori, T., Uemura, Y., Nagatsu, I., Kikuchi, H., Mizuno, N., 1989. Tyrosine hydroxylase-like immunoreactive neurons in the striatum of the rat. Neurosci Lett 97, 6–10.

Tay, T.L., Ronneberger, O., Ryu, S., Nitschke, R., Driever, W., 2011. Comprehensive catecholaminergic projectome analysis reveals single-neuron integration of zebrafish ascending and descending dopaminergic systems. Nature communications 2, 171.

Terzibasi, E., Lefrancois, C., Domenici, P., Hartmann, N., Graf, M., Cellerino, A., 2009. Effects of dietary restriction on mortality and age-related phenotypes in the short-lived fish Nothobranchius furzeri. Aging Cell 8, 88–99.

Valenzano, D.R., Terzibasi, E., Genade, T., Cattaneo, A., Domenici, L., Cellerino, A., 2006. Resveratrol prolongs lifespan and retards the onset of age-related markers in a short-lived vertebrate. Curr Biol 16, 296–300.

Vigh, B., Manzano, M.J., Zadori, A., Frank, C.L., Lukats, A., Rohlich, P., Szel, A., David, C., 2002. Nonvisual photoreceptors of the deep brain, pineal organs and retina. Histol Histopathol 17, 555–590.

von, E.U., 1961. Occurrence and distribution of catecholamines in the fish brain. Acta Physiol Scand 52, 62–64.

Wang, Y., Liu, W., Yang, J., Wang, F., Sima, Y., Zhong, Z.M., Wang, H., Hu, L.F., Liu, C.F., 2017. Parkinson’s disease-like motor and non-motor symptoms in rotenone-treated zebrafish. Neurotoxicology 58, 103–109.

Wicht, H., Northcutt, R.G., 1994. An immunohistochemical study of the telencephalon and the diencephalon in a Myxinoid jawless fish, the Pacific hagfish, Eptatretus stouti. Brain Behav Evol 43, 140–161.

Wullimann, M.F., Rupp, B., Reichert, H., 1996. Neuroanatomy of the Zebrafish Brain: A Topological Atlas. Basel: Birkhäuser.

Xavier, A.L., Fontaine, R., Bloch, S., Affaticati, P., Jenett, A., Demarque, M., Vernier, P., Yamamoto, K., 2017. Comparative analysis of monoaminergic cerebrospinal fluid-contacting cells in Osteichthyes (bony vertebrates). The Journal of comparative neurology 525, 2265–2283.

Yamamoto, K., Ruuskanen, J.O., Wullimann, M.F., Vernier, P., 2010. Two tyrosine hydroxylase genes in vertebrates New dopaminergic territories revealed in the zebrafish brain. Mol Cell Neurosci 43, 394–402.

Yamamoto, K., Vernier, P., 2011. The evolution of dopamine systems in chordates. Front Neuroanat 5, 21.

