## Supplementary figures for "Organization of the catecholaminergic system in the short-lived fish *Nothobranchius furzeri*"

### *Supplementary Material*

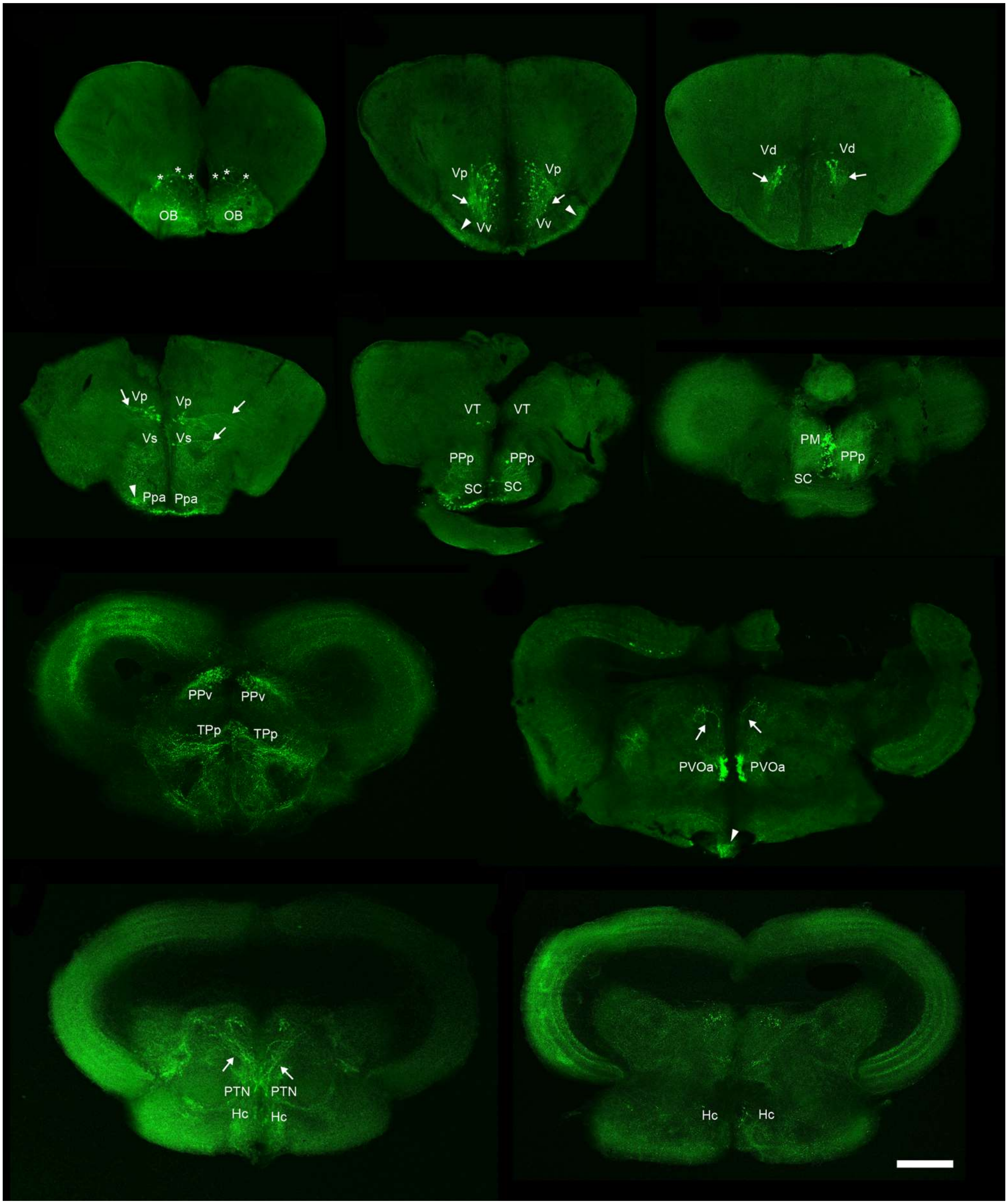

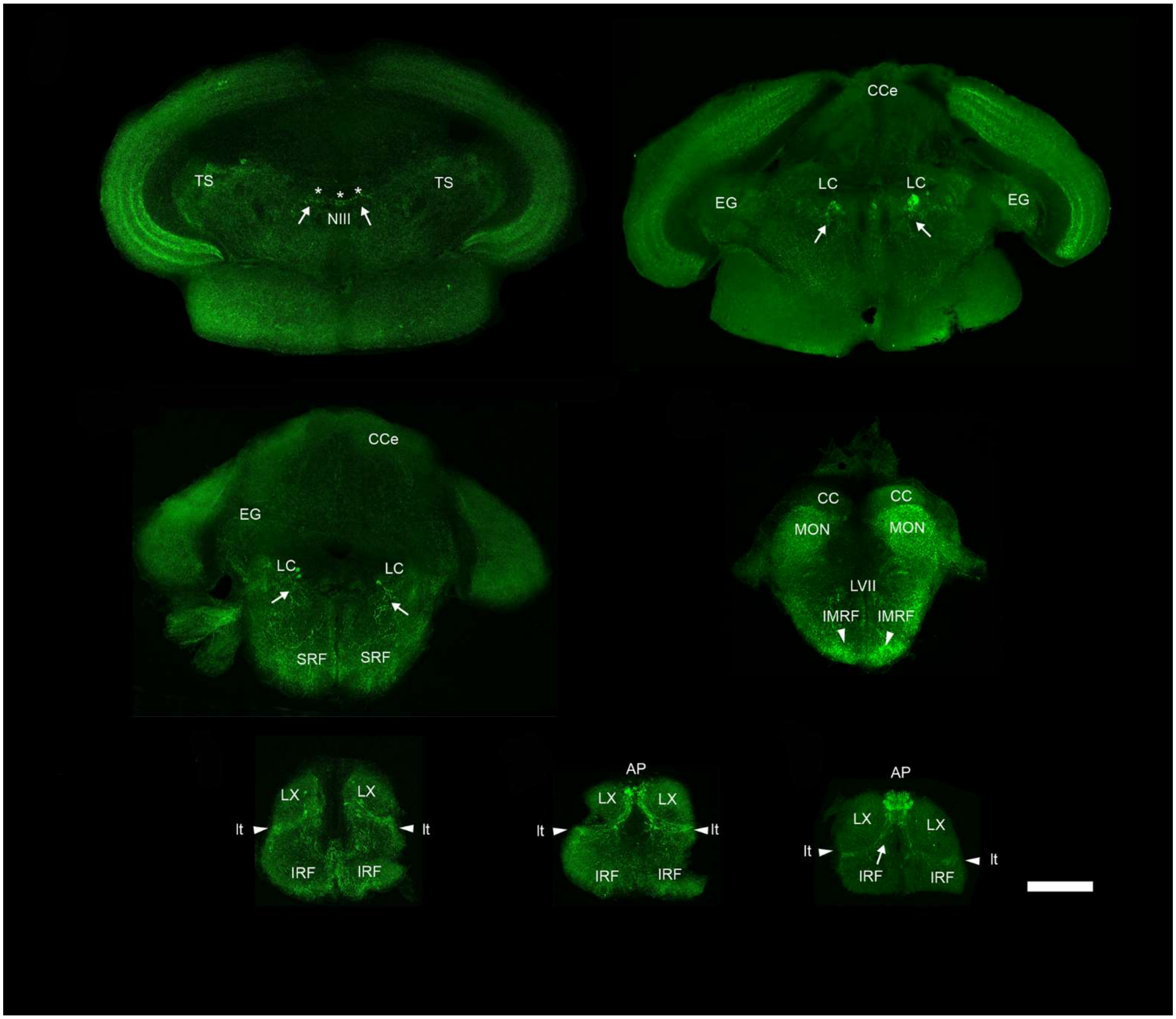


**Supplementary Figure 1. Photomicrographs of transverse sections (in rostral to caudal order) showing the anti-TH immunofluorescence in the brain of *Nothobranchius furzeri*.** Asterisks point neuronal somas, arrows show the direction of the processes and arrowheads indicate regions with intense density of TH+ neuropil or tracts. For more details see the text and for abbreviations see list. Scale bar, 500 µm.

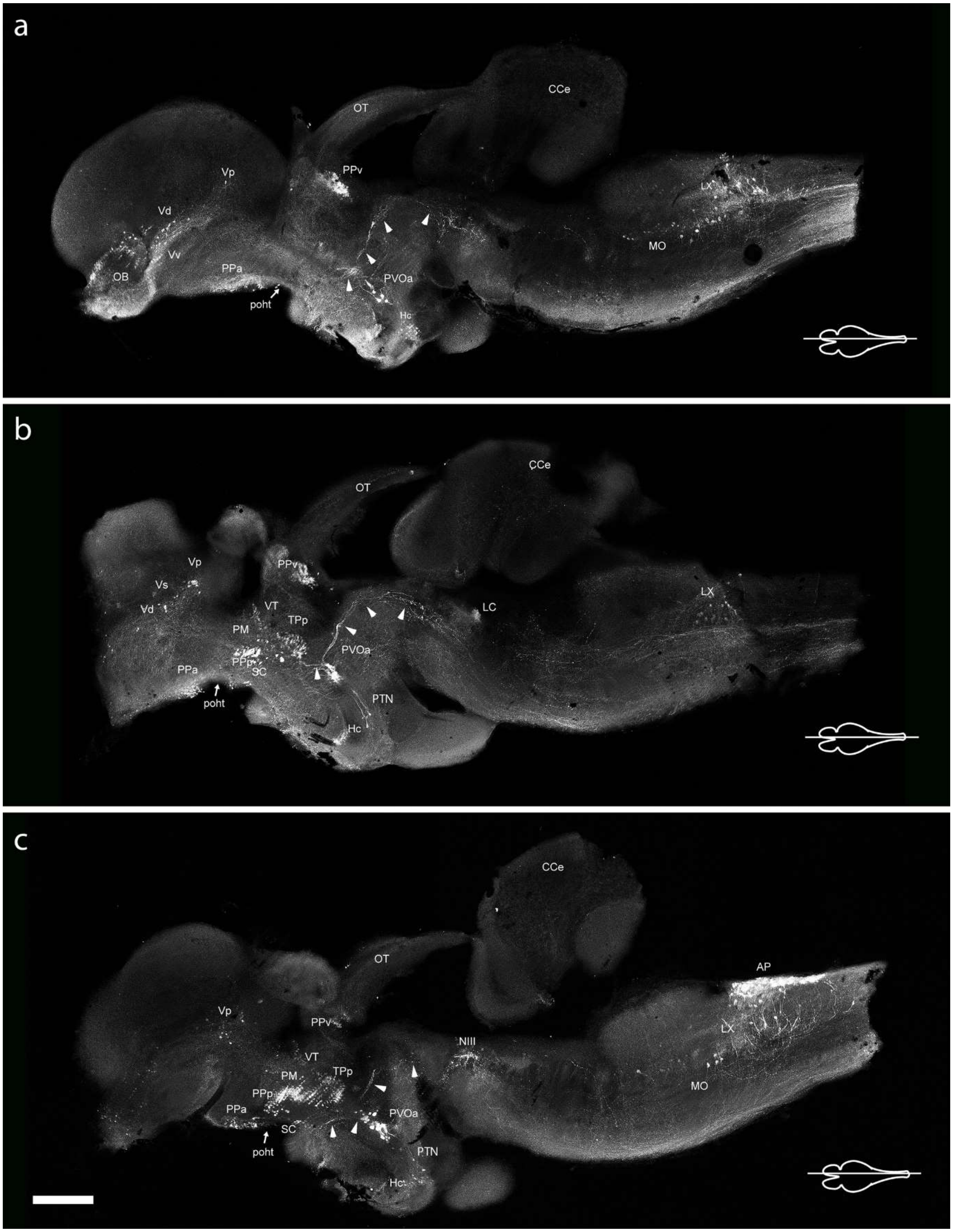


**Supplementary Figure 2. Photomicrographs of para-sagittal sections near the midline (from lateral towards the midline), showing the distribution of TH+ neuronal groups and fibers/tracts through the brain of *Nothobranchius furzeri*.** Arrowheads point to TH+ fibers arising from the paraventricular organ-accompanying cells and arrows indicate the preopticohypothalamic tract. For more details see the text and for abbreviations see list. Scale bar, 500 µm.

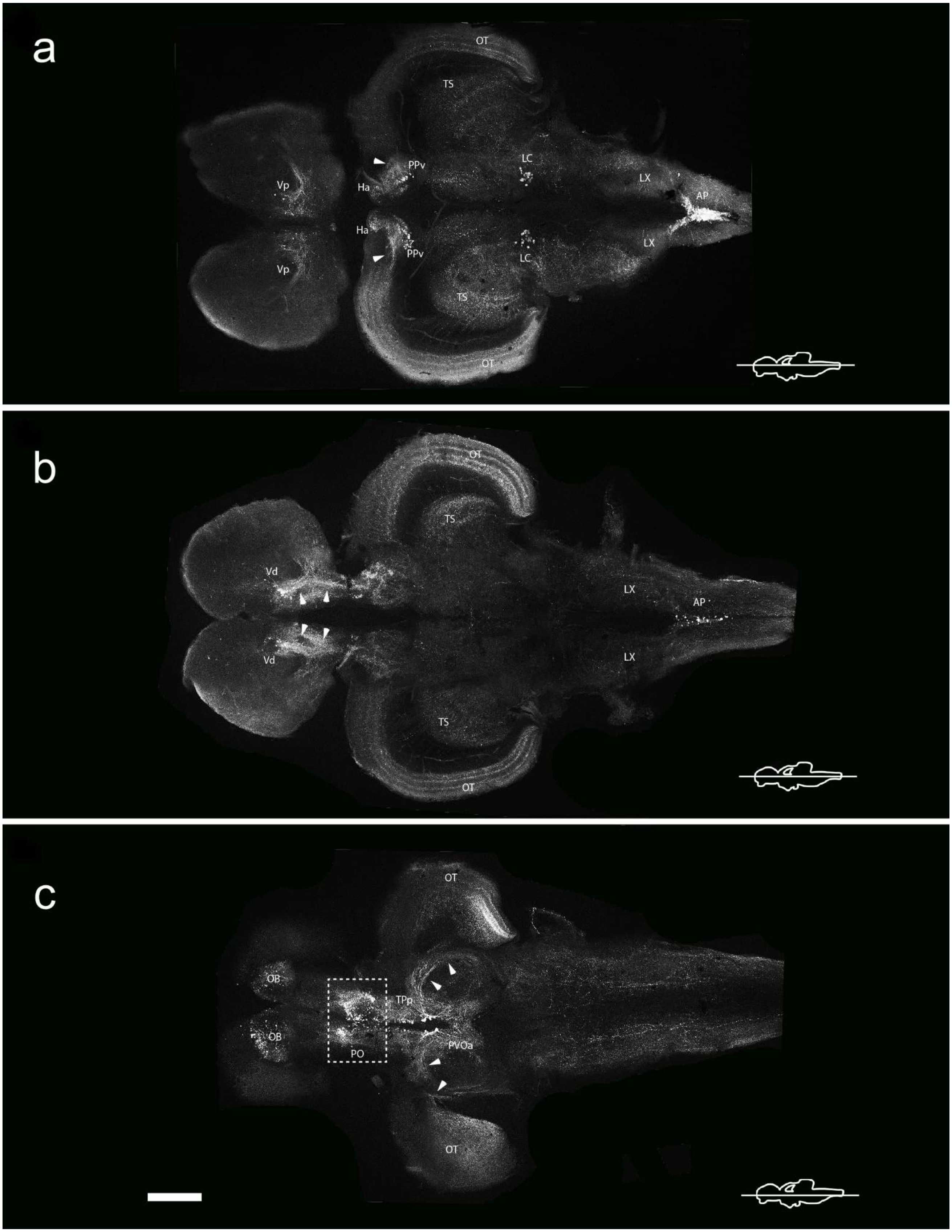
a
OT
TS
PPv
LC
Vp
Ha
LX
AP
Ha
PPv
Vp
LC
LX
TS
OT
b
OT
TS
Vd
LX
AP
Vd
LX
TS
OT
c
OT
OB
TPp
OB
PVOa
PO
OT

**Supplementary Figure 3. Photomicrographs of horizontal sections (in dorsal to ventral order) showing the distribution of TH+ cellular groups and fibers/tracts along the rostrocaudal axis of the brain in *Nothobranchius furzeri*.** Arrowheads in **(A)** indicate the TH+ fibers of the ventral periventricular pretectal nucleus reaching the optic tectum, in **(B)** the fibers from the posterior zone of ventral telencephalon following a caudal direction, and in **(C)** the direction of processes from the paraventricular organ-accompanying cells. For more details see the text and for abbreviations see list. Scale bar, 500 μm.
